# Calretinin positive neurons form an excitatory amplifier network in the spinal cord dorsal horn

**DOI:** 10.1101/673533

**Authors:** KM Smith, TJ Browne, O Davis, A Coyle, KA Boyle, M Watanabe, SA Dickinson, JA Iredale, MA Gradwell, P Jobling, RJ Callister, CV Dayas, DI Hughes, BA Graham

## Abstract

The passage of nociceptive information is relayed through the spinal cord dorsal horn, a critical area in sensory processing. The neuronal circuits in this region that underpin sensory perception must be clarified to better understand how dysfunction can lead to pathological pain. This study used an optogenetic approach to selectively activate neurons that contain the calcium-binding protein calretinin (CR). We show that CR^+^ interneurons form an interconnected network that can initiate and sustain enhanced excitatory signaling, and directly relays signals to lamina I projection neurons. *In vivo* photoactivation of CR^+^ interneurons resulted in a significant nocifensive behavior that was morphine sensitive and cause a conditioned place aversion. Furthermore, halorhodopsin-mediated inhibition of CR^+^ interneurons elevated sensory thresholds. These results suggest that neuronal circuits in the superficial dorsal horn that involve excitatory CR^+^ neurons are important for the generation and amplification of pain, and identify these interneurons as a future analgesic target.

## Introduction

All sensory information from the body, including nociception, is first relayed into the spinal cord dorsal horn (DH), where this afferent input can be modulated, gated and prioritized before being relayed to higher centers for sensory perception (Todd, 2010, Peirs and Seal, 2016). It is well established that alterations to neuronal circuits within the DH can directly contribute to neuropathic and inflammatory pain, as well as persistent itch (Basbaum et al., 2009, Braz et al., 2014). Despite this being a region of immense biological importance, our understanding of the neuronal circuits associated with particular sensory modalities remains limited (Todd, 2010, Peirs and Seal, 2016). To address this knowledge gap, several groups have recently implicated neurochemically-distinct subpopulations of DH interneurons with the perception of both acute and chronic pain states (Smith et al., 2015, Peirs et al., 2015, Petitjean et al., 2015, Duan et al., 2014).

Historically, much of the research effort on DH circuits has focused on inhibition (Zeilhofer et al., 2012), and a growing number of discrete inhibitory interneuron populations have now been identified as substrates for sensory gating in the spinal cord (Petitjean et al., 2015, Duan et al., 2014, Foster et al., 2015, Cui et al., 2016). In contrast, our understanding of the role excitatory interneurons play in sensory processing is far less developed. Generally, excitatory DH populations are considered to provide polysynaptic relays linking circuits dedicated to innocuous and noxious sensory input, with inhibitory populations normally modulating the passage of information through these pathways (Duan et al., 2014, Takazawa and MacDermott, 2010, Punnakkal et al., 2014). Such a limited role for excitatory interneurons is surprising given they outnumber inhibitory interneurons by 2:1 in superficial laminae (Polgar et al., 2013), suggestive of a more complex role. Furthermore, detailed paired recording studies have also shown that ∼85% of the synaptic connections in the DH are excitatory (Santos et al., 2007).

Notably, we have recently shown that most calretinin-expressing (CR) neurons in laminae I and II exhibit specific electrophysiological, morphological and neurochemical properties consistent with an excitatory phenotype and respond to noxious peripheral stimulation (Smith et al., 2015, Smith et al., 2016). In fact, chemogenetic activation of CR^+^ neurons has been shown to cause nocifensive behaviors and DH activation patterns consistent with mechanical hypersensitivity (Peirs et al., 2015), whereas genetic ablation of CR^+^ neurons can cause a selective loss of light punctate touch sensation (Duan et al., 2014). Prior work has established that a specific excitatory interneuron population in the deep dorsal horn that transiently express VGLUT3 relays low threshold input to CR^+^ neurons (Peirs et al., 2015), however, a major limitation remains the lack of detailed information on the postsynaptic circuits engaged by the CR^+^ population to drive behavioral responses.

Here, we take an optogenetic approach to resolve the neuronal circuits excited by CR^+^ neurons in laminae I and II, and determine the functional significance of these neurons for sensory processing and perception. The postsynaptic targets of CR^+^ neurons were identified combining optogenetic stimulation with *in vitro* electrophysiology, and also producing activation maps in anesthetized animals. This identified somatostatin^+^ neurons, neurokinin 1 receptor positive spinoparabrachial projection neurons, and CR^+^ neurons themselves among recipient populations for CR^+^ input. Together, these populations form a highly integrated excitatory network that is able to amplify dorsal horn circuit activity including downstream neural targets. Using *in vivo* optogenetic stimulation in awake and behaving animals we were also able to show that spinal activation of CR^+^ neurons induces nocifensive behavior.

## Results

### Optogenetic activation of spinal CR^+^ neurons

To study spinal CR^+^ neuron connectivity and function in sensory processing, CR-Cre mice (Cr-IRES-Cre) were crossed with loxP-flanked-ChR2-eYFP mice (Ai32) to generate offspring where ChR2 was expressed in CR^+^ neurons (CR^cre^;Ai32). These mice exhibited characteristic ChR2-eYFP expression in neurons and fibers located in the superficial DH of the spinal cord forming a plexus that was concentrated in lamina IIo (Supplementary Figure 1A). This is consistent with the known pattern of CR expression in this spinal cord region(Lu and Perl, 2003). Comparison with immunolabelling for CR confirmed ChR2-eYFP expression was highly localized to the CR^+^ population with 78.3 ± 4 % (St. Dev.) of CR^+^ neurons expressing ChR2-eYFP (1454 cells counted in 3 animals), and 71.5% ± 2% of ChR2-eYFP^+^ neurons expressing CR (1767 cells counted in 3 animals). Consistent with our previous work, most CR neurons exhibited characteristic electrophysiological features indicative of excitatory interneurons (Supplementary Figure 1B). In voltage clamp, these ChR2-eYFP expressing cells exhibited robust inward photocurrents in response to photostimulation (n = 29 cells from 16 animals), which increased with stimulation intensity (0.01-16 mW, Supplementary Figure 1C). In current clamp, photostimulation evoked AP discharge, and the ChR2-eYFP neurons were able to reliably follow repetitive stimulation trains up to 10 Hz, however, reliability decreased at higher frequencies (Supplementary Figure 1D). The latency between photostimulation onset and AP discharge (i.e. recruitment delay) across this sample was 3.29 ± 0.21 ms. We also assessed whether the subset of CR^+^ neurons, identified in our previous work as inhibitory interneurons (*Atypical* CR^+^ neurons) expressed ChR2-eYFP (n = 13 cells from 9 animals). These cells exhibit morphological and electrophysiologial features consistent with an inhibitory phenotype (Supplementary Figure 2A). Photostimulation in this inhibitory subset of ChR2-eYFP expressing neurons evoked larger inward photocurrents than observed in the excitatory population (459.72 ± 34.85 pA *vs*. 233.66 ± 56.16 pA), which similarly increased with photostimulation intensity (Supplementary Figure 2B). The inhibitory ChR2-eYFP population could also reliably follow repetitive photostimulation at rates up to 10Hz, but had a shorter recruitment time than excitatory ChR2-eYFP neurons (2.39 ± 0.21 ms *vs.* 3.29 ± 0.38 ms, Supplementary Figure 2C). Together, these data indicate the CR^cre^;Ai32 mouse provides optogenetic control of both excitatory and inhibitory CR^+^ populations.

### CR-ChR2-activated microcircuits

Channelrhodopsin-2 assisted circuit mapping (CRACM) was used to study the connectivity of CR-ChR2 neurons within DH microcircuits. Brief full-field photostimulation (16 mW, 1 ms) was applied to assess excitatory postsynaptic responses across various DH populations (n = 73 cells from 27 animals). Strikingly, robust synaptic responses were observed in the CR-ChR2 neurons themselves (Figure 1B). Specifically, photostimulation of CR^+^ neurons produced responses that included an immediate photocurrent and short latency optically evoked excitatory postsynaptic currents (oEPSCs) that were blocked by bath applied CNQX (10 μM). In order to analyse the oEPSCs, pharmacologically isolated photocurrents (after CNQX) were first subtracted from the original response, separating oEPSCs (Supplementary Figure 3A). oEPSCs were observed in 96.5% of these recordings (28/29) indicating a high degree of interconnectivity in the CR-ChR2 population. A defined a window for direct connection latencies was characterised by adding a delay of 2.5 ms (taken from previous paired recording studies(Santos et al., 2007, Lu and Perl, 2003)) to the average AP recruitment delay for excitatory CR^+^ neurons (3.29 ± 0.38 ms, Supplementary Figure 1), allowing for AP conduction and synaptic delay. The distribution of oEPSC latencies in CR-ChR2 neurons suggested they receive both a direct and delayed input following photostimulation (35% direct, 65% delayed, Supplementary Figure 4A).

**Figure 1.**
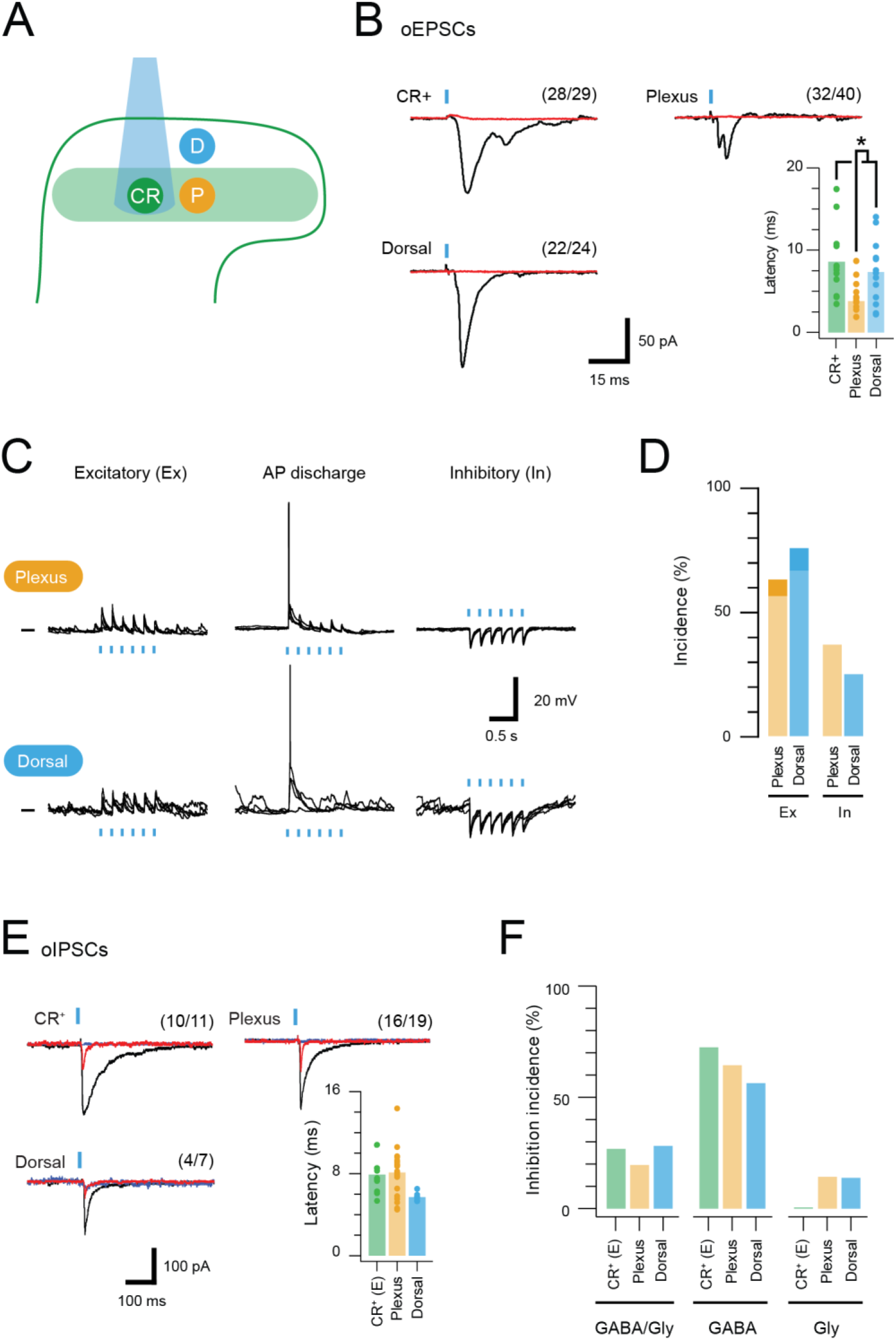
CR-ChR2 neurons provide excitatory drive throughout the DH. **(A)**, Schematic shows DH populations assessed for CR-ChR2-evoked excitatory input: CR-ChR2^+^ neuron (green), interneurons (yellow) located within the CR^+^ plexus (light green shading), and interneurons located dorsal to the CR^+^ plexus (blue). **(B)**, Photostimulation (16 mW, 1 ms) evoked robust inward currents under voltage clamp in each DH population. Traces show averaged response (black) to photostimulus (blue bar), CNQX (10 μM) abolished all responses (red). Values on each trace show number of recordings that exhibited a light induced inward current. Bar graph shows group data comparing oEPCS latency, which was significantly shorter in interneurons within the CR plexus (p=0.047). **(C)**, Representative traces show responses during photostimulation recorded from interneurons within the CR^+^ plexus (upper) and dorsal to this region (lower), in current clamp. In some neurons photostimulation only caused subthreshold depolarization (excitatory, left); in others depolarization evoked AP discharge (center), while in some neurons the postsynaptic response during photostimulation was inhibitory in the form of transient membrane hyperpolarisations (right). Photostimulation applied at a membrane potential of −60mV. **(D)**, Bar graphs show group data on the incidence of photostimulation responses. Darker shading denotes percentage of excitatory responses that cause AP discharge in each group. **(E)**, Traces show averaged optically evoked inhibitory postsynaptic currents (oIPSCs) recorded in response to photostimulation (black trace), and following bath applied bicuculline (10µM, red trace) and strychnine (1µM, blue trace), left to right. Bar graph compares the latency of inhibitory responses from photostimulation onset. **(C)**, Photostimulation-evoked inhibitory responses were classed as mixed (GABA/glycine, left), GABA-dominant (middle) or glycine dominant (right) based on bicuculline sensitivity. The incidence of each form of inhibition was similar across the populations assessed.

CRACM was also applied while recording from neurons lacking ChR2 both within or dorsal to the CR^+^ plexus (LII_o_), showing that both plexus (32/40) and dorsal (22/24) populations received CR-ChR2 neuron input (Figure 1B). Using the same defined window for direct and delayed input, plexus recordings received mostly direct input (75% direct, 25% delayed, Supplementary Figure 4A), whereas recordings dorsal to the CR^+^ plexus exhibited a similar level of direct and delayed oEPSC input (57% direct, 43% delayed, Supplementary Figure 4A). Comparison of oEPSC characteristics identified a significantly shorter onset of the oEPSC response (Figure 1B) for neurons within the CR^+^ plexus compared to other populations (plexus = 4.75 ± 0.59 ms *vs*. CR-ChR2 = 8.61 ± 1.23 ms, p=0.012; dorsal = 7.34 ± 1.06 ms, p=0.047). In contrast, oEPSC time-course was similar across recordings (Table 1; rise time: CR-ChR2 = 2.68 ± 0.41 ms; plexus = 2.89 ± 0.63 ms; and dorsal = 3.22 ± 0.64 ms. Half Width: CR-ChR2 = 4.90 ± 0.62 ms; plexus = 5.40 ± 1.60 ms; and dorsal = 6.61 ± 1.16 ms). These features combined to generate similar oEPSC charge across the sampled populations (CR-ChR2 = 0.66 ± 0.20 pA.s; plexus = 0.52 ± 0.17 pA.s; dorsal = 1.93 ± 0.98 pA.s). Thus, activation of CR-ChR2 neurons produces excitation that unsurprisingly arrives first on nearby populations within the ChR2-eYFP plexus, before it reaches neurons dorsal to this region. In addition, interconnectivity of CR-ChR2 neurons indicates they form an excitatory network likely to enhance activity within the DH when recruited.

**Table 1:** Photostimulation response characteristics.

| Neuron type | Input | n | Latency (ms) | Amplitude (pA) | Rise time (ms) | Half-width (ms) | Charge (pA.ms) |
| --- | --- | --- | --- | --- | --- | --- | --- |
| CR-ChR2 (CR) | oEPSC | 12 | 8.6 ± 1.2 | 87.59 ± 20.04 | 2.67 ± 0.41 | 4.90 ± 0.62 | 0.66 ± 0.20 |
| Plexus (P) | oEPSC | 11 | 4.8 ± 0.6 *<sup>CR, D, PN</sup> | 51.42 ± 19.88 | 2.89 ± 0.63 | 5.40 ± 1.60 | 0.52 ± 0.17 |
| Dorsal (D) | oEPSC | 13 | 7.3 ± 1.1 | 126.72 ± 52.03 | 3.22 ± 0.64 | 6.61 ± 1.16 | 1.93 ± 0.98 |
| PN (PN) | oEPSC | 13 | 7.9 ± 1.4 | 70.68 ± 131.40 | 5.54 ± 1.44 *<sup>CR, P, D</sup> | 10.82 ± 1.60 *<sup>CR, P, D</sup> | 1.35 ± 0.55 |
| CR-ChR2 (CR) | Mixed-oIPSC (M) | 10 | 7.91 ± 0.53 | 146.43 ± 70.33 | 14.82 ± 3.90 | 74.39 ± 26.28 | 29.25 ± 23.87 |
|  | Gly-oIPSC (G) | 10 |  | 52.66 ± 12.75 | 4.36 ± 1.17 *<sup>M</sup> | 16.72 ± 3.42 *<sup>M</sup> | 9.77 ± 7.26 |
| Plexus (P) | Mixed-oIPSC (M) | 16 | 8.32 ± 0.61 | 237.16 ± 70.72 | 7.74 ± 0.60 | 71.70 ± 10.00 | 27.29 ± 9.30 |
|  | Gly-oIPSC (G) | 16 |  | 104.28 ± 40.55 *<sup>M</sup> | 6.27 ± 1.33 | 40.56 ± 19.30 *<sup>M</sup> | 8.17 ± 4.13 *<sup>M</sup> |
| Dorsal (D) | Mixed-oIPSC (M) | 4 | 5.72 ± 0.53 | 224.87 ± 76.67 | 5.99 ± 1.67 | 19.75 ± 2.93 | 17.95 ± 10.66 |
|  | Gly-oIPSC (G) | 4 |  | 81.34 ± 50.52 *<sup>M</sup> | 2.645 ± 0.35 *<sup>M</sup> | 15.87 ± 5.58 | 5.18 ± 3.46 *<sup>M</sup> |

Values are mean ± SEM. \* denotes p<0.05 between cell types (CR vs. P vs. D vs. PN), or oIPSC type (M vs. G)

The impact of CR-ChR2 photostimulation on the activity of postsynaptic populations was also assessed in current clamp (n = 22 cells from 12 animals). Three response types were typically distinguished in these recordings; i) subthreshold excitatory responses, ii) suprathreshold excitatory responses (i.e. evoked AP discharge), and iii) inhibitory responses (Figure 1C). Responses were assessed in neurons within the ChR2-YFP plexus (and dorsal to this region, but not CR-ChR2 neurons, as they were directly activated to photostimulation. The incidence of each responses was similar among the ChR2-eYFP plexus and dorsal recordings (Figure 1D) including excitatory (56.3% and 66.6%) and inhibitory (37.5% and 25%) responses, with few neurons responding with AP discharge (6.2% and 8.4%).

Given the appearance of inhibitory responses in the above recordings, and the likelihood that inhibitory CR-ChR2 neurons were also activated by photostimulation, CRACM also assessed inhibitory connections within the dorsal horn (n = 29 cells from 13 animals). Optically evoked inhibitory postsynaptic currents (oIPSCs) were observed in all neuron populations studied (Figure 1E – CR-ChR2 10/11, plexus 16/19, and dorsal neurons 4/7). Comparison of oIPSC characteristics showed that oIPSC latency was similar among these recordings (CR-ChR2 = 7.9 ± 0.5 ms, and dorsal = 5.7 ± 0.1 ms). To determine the contribution of direct and delayed circuits to this response, a latency window for oIPSC components to be considered direct was calculated (as above for oEPSCs) using the inhibitory CR-ChR2 neuron recruitment latency of 2.39 ± 0.21 ms, and 2.5 ms to account for AP conduction and synaptic delay(Santos et al., 2007, Lu and Perl, 2003). All neuron types exhibited responses consistent with direct and delayed oIPSC components (Supplementary Figure 4B). Delayed oIPSCs components dominated in neurons within the CR^+^ plexus (80.5% delayed *vs*. 19.5% direct), whereas a similar mix of direct and delayed oIPSC components were recorded in excitatory CR-ChR2 neurons and neurons dorsal to the CR^+^ plexus (excitatory CR-ChR2 = 53% delayed *vs*. 47% direct, and dorsal 59% delayed *vs*. 41% direct). Other oIPSC properties were generally similar across neuron types (Table 1). Importantly, as both GABA and glycine can mediate fast synaptic inhibition in the spinal DH, sequential pharmacology was used to differentiate these neurotransmitters. Photostimulation-evoked oIPSC responses were isolated by bath application of CNQX (10 μM) and then GABAergic oIPSC components were blocked with bicuculline (10 μM), before any remaining oIPSCs were abolished with strychnine (1 μM). Comparison of oIPSCs recorded before and after bicuculline block assessed the contribution of GABA and glycine to these photostimulation responses. In this way, an oIPSC amplitude decrease of 80% or greater in bicuculline indicated GABA-dominant input, whereas a decrease of less than 20% in bicuculline indicated a glycine-dominant input. oIPSCs with intermediate bicuculline-sensitivity were classified as mixed (i.e. both GABAergic and glycinergic). Across all recordings, GABA-dominant responses were most common (Figure 1F: excitatory-CR-ChR2 = 72%, plexus = 65%, dorsal = 58%). Glycine dominant responses were rare, and not observed at all in excitatory CR-ChR2 neurons, with remaining cells receiving mixed inhibition. Together, these data show that in addition to a range of excitatory circuits recruited by CR-ChR2 photostimulation, a widely distributed pattern of inhibition is also activated by CR-ChR2 neuron recruitment. Short latency direct inhibition likely comes through direct photostimulation of inhibitory CR-ChR2 neurons, whereas polysynaptic pathways recruited by photostimulation of excitatory CR-ChR2 neurons are best placed to produce longer latency indirect inhibitory responses.

### Plasticity in the CR-ChR2 network

The interconnectivity of the CR-ChR2^+^ population and multicomponent responses to brief photostimulation (direct and delayed) suggested these neurons might be capable of producing sustained activation within DH circuits. To test this hypothesis, photostimulation duration was extended (10 s @ 10 Hz, 10 ms pulses at 16 mW) and spontaneous EPSC (sEPSC) frequency before and immediately following photostimulation were compared (Figure 2). Recordings in spinal slices from CR^cre^;Ai32 animals (n = 4) targeted CR-ChR2^+^ neurons due to their coupling and predominantly excitatory phenotype, but also sampled other unidentified DH neurons, and some inhibitory CR-ChR2^+^ neurons (differentiated from the excitatory CR^+^ population by their discharge characteristics). These recordings exhibited a range of pre-stimulation and post-stimulation sEPSC frequency relationships, however, post-stimulation sEPSC frequency was dramatically increased in a subset of CR-ChR2^+^ neurons (Figure 2A-B). A threshold of 4 standard deviations above the mean pre-stimulation sEPSC frequency was set to confidently identify recordings with increased post-photostimulation sEPSC frequency. Using this criterion one third of excitatory CR-ChR2^+^ recordings (4/12) exhibited increased post-stimulation sEPSC frequency (Figure 2B). In contrast, poststimulation sEPSC frequency did not increase in unidentified DH neurons (0/9), or inhibitory CR-ChR2^+^ neurons (0/5). While these potentiated responses could result from the specific connectivity patterns in the CR^+^ network, they may also relate to direct activation of photocurrents in these neurons, or the magnitude of evoked oEPSC during photostimulation. Despite this, there was no correlation between the degree of potentiation and either photocurrent amplitude (Figure 2C left; r^2^= 0.00002) or oEPSC amplitude (Figure 2C right; r^2^= 0.056). Peristimulus histograms (Figure 2D) compared CR-ChR2^+^ neuron responses that exhibited increased post-stimulation sEPSC frequency (n=4) with CR-ChR2^+^ neurons that exhibited similar baseline sEPSC frequency but no poststimulation increase (n=5). This highlighted the dramatic and prolonged nature of enhanced excitatory synaptic activity in the post-stimulation period, taking ∼20 seconds before returning to baseline. Together, these results are compatible with a model that features feedback excitation within the CR^+^ network, capable of maintaining elevated excitatory signalling beyond the initial excitatory stimulus.

**Figure 2.**
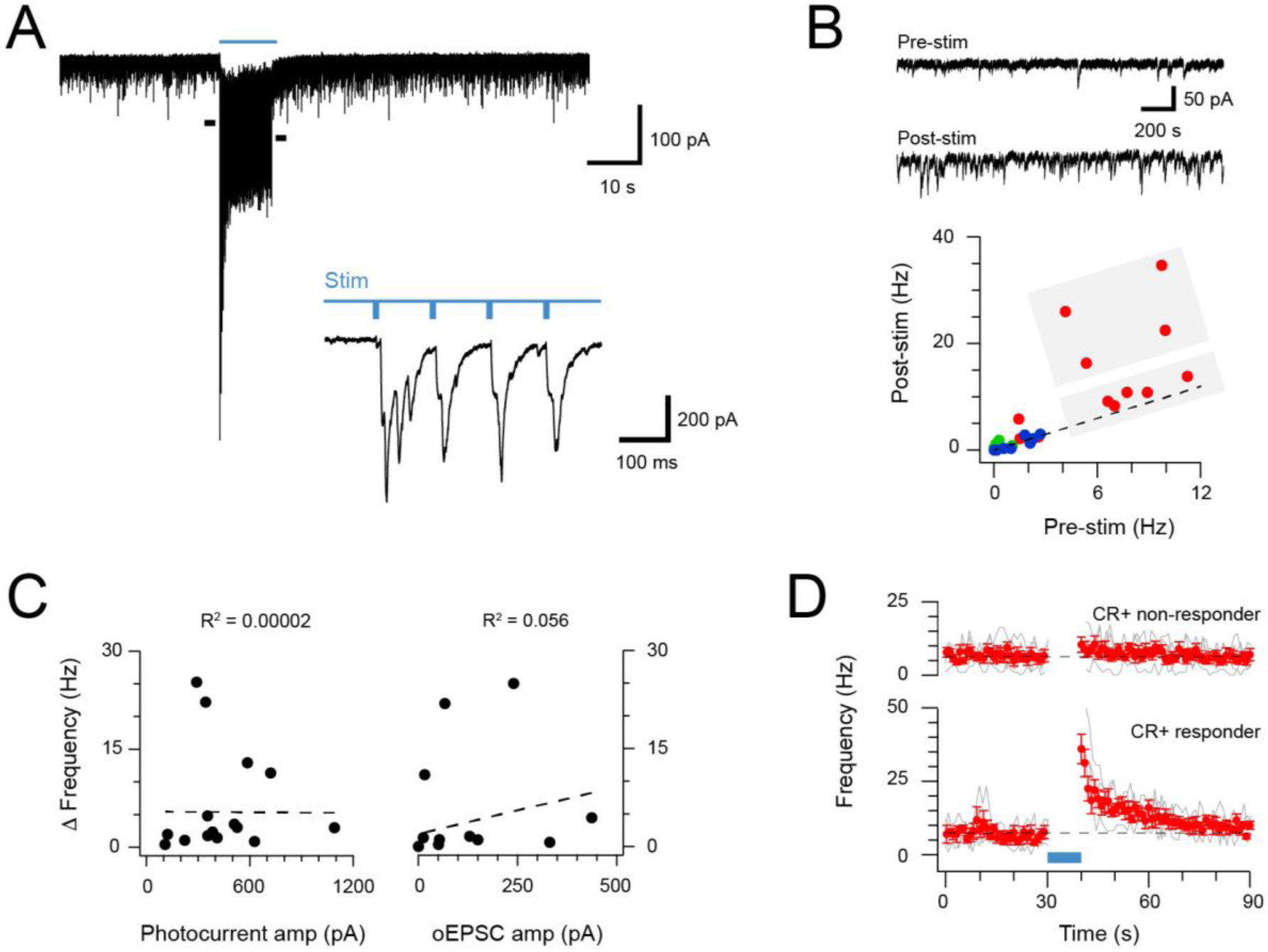
Extended CR-ChR2 photostimlation enhances spontaneous excitatory activity. **(A)**, spinal cord slice recordings from a CR-ChR2^+^ neuron showing spontaneous excitatory postsynaptic currents before, during and following full field photostimulation (blue bar, 16 mW, 10 ms pulses @ 10 Hz, 10 s). Inset shows onset of photostimulation and response on expanded time scale. Note a dramatic increase in sEPSC frequency persists following the photostimulation period. **(B)**, Traces (upper) show 2 s pre and post photostimlation on an expanded timescale from **A**. Plot (lower) shows group data comparing sEPSC frequency in the pre- and post-photostimulation (excitatory CR-ChR2 cells = red, inhibitory CR-ChR2 cells = green, unidentified DH cells = blue). Data on or near the unity line (dashed) indicates little change between pre- and post-photostimulation sEPSC frequency, however, four CR-ChR2 cells exhibited a substantial increase in post photostimulation sEPSC frequency (large grey box). **(C)** Plots compare pre- to post-photostimulation frequency sEPSC (Δ frequency) with photocurrent and photostimulated oEPSC amplitudes (left and right, respectively). There was no correlation between Δ frequency and either property. **(D)** Plots compare mean sEPSC frequency (red) across photostimulation protocol for CR-ChR2 cells deemed to exhibit a post-photostimulation increase (n=4, post-photostimulation sEPSC frequency exceeded mean pre-photostimulation sEPSC frequency ± 4SD), and CR-ChR2 cells with a similar baseline sEPSC frequency, but no post-photostimulation change.

### Distinct DH populations are activated by CR-ChR2 neurons

To identify the DH populations postsynaptic to CR-ChR2 neurons, a deeply anesthetized preparation was used and photostimulation (10 mW, 10 ms @ 10 Hz for 10 min) was delivered to the exposed dorsal surface of the spinal cord in CR^Cre^;Ai32 mice. Spinal cords were subsequently processed and immunolabeled for the activity marker Fos, YFP, and neurochemical markers commonly used to differentiate DH populations implicated in pain pathways(Peirs and Seal, 2016, Duan et al., 2014). Robust Fos-protein induction was restricted to the photostimulation area (Figure 3A-B). Importantly, no Fos-positive profiles were found in control animals (identical photostimulation in CReGFP mice, n = 3 animals) confirming the specificity of photostimulation evoked Fos expression in CR^Cre^;Ai32 mice. Of the Fos^+^ profiles, approximately one third expressed YFP (21.2 ± 4.1 of 73.8 ± 9.8 neurons, 12 animals) indicating these neurons expressed ChR2 and were directly activated by photostimulation. The remaining two thirds of Fos^+^ neurons represented postsynaptic targets of the CR-ChR2^+^ population. Approximately 10% of these cells were NK1R-expressing lamina I neurons (Figure 3C, 4.7 ± 4.6 of 41.3 ± 2.51 neurons; 3 animals). As immunolabelling for NK1R was confined to the cell membrane and showed no evidence of internalisation, we conclude that activation of these putative projection neurons resulted from glutamatergic synaptic input derived from photostimulated CR-ChR2^+^ spinal interneurons or their postsynaptic targets.

**Figure 3.**
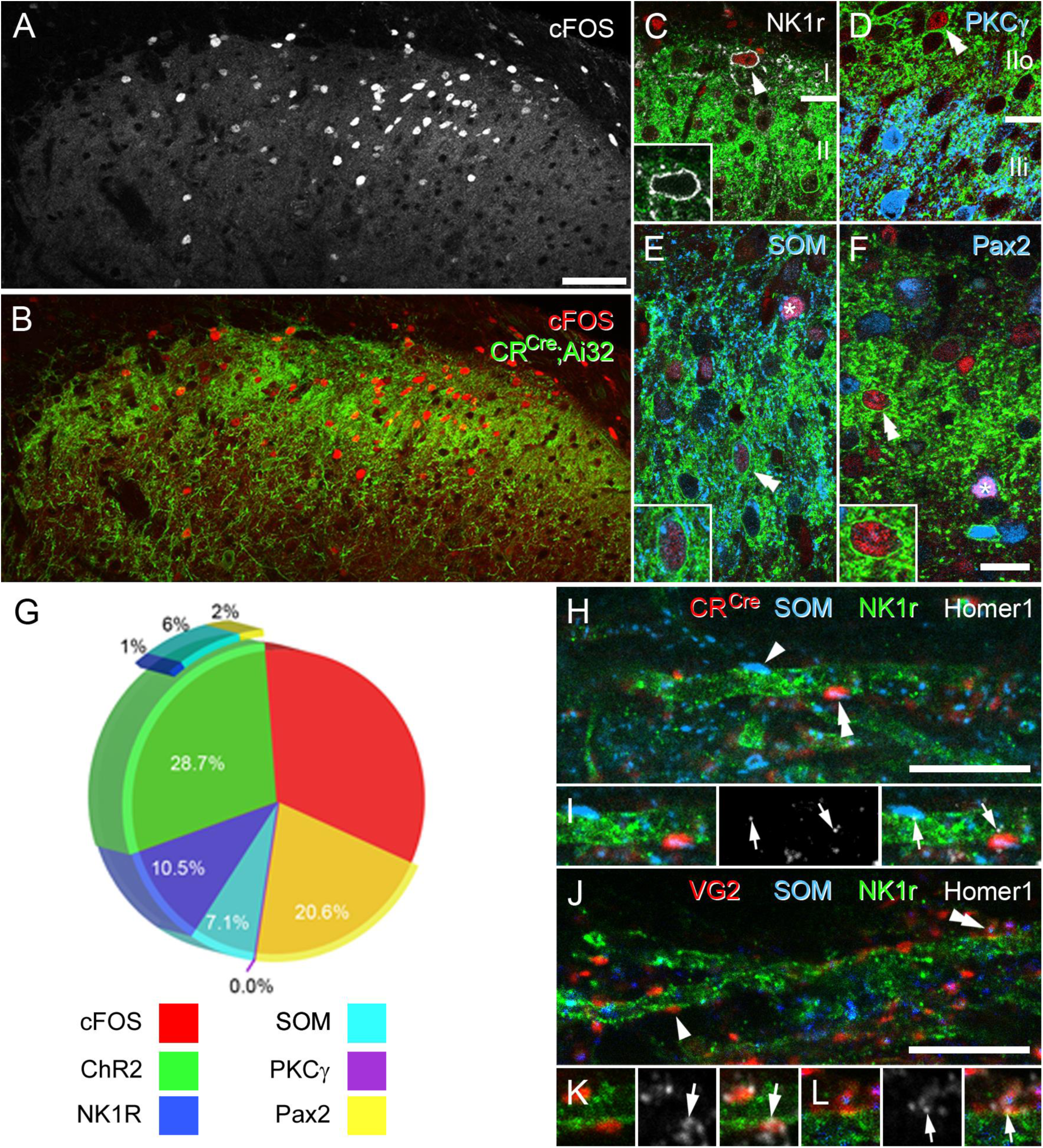
CR-ChR2 neuron photostimulation activates multiple DH neuron populations. **(A)**, Following photostimulation in a deeply anaesthetized CR^cre^;Ai32 mouse, robust cFos-IR profiles (white) were detected in laminae I and II primarily. **(B)**, These cFOS-IR cells (red) were restricted to the ipsilateral DH, and largely confined to the CR-ChR2-YFP plexus (green). **(C)**, Lamina I neurons often expressed cFOS, and these were commonly immunolabelled for NK1R (arrowhead; white). In these cells, NK1R-immunolabelling was confined to the cell membrane (inset). **(D)**, Immunolabelling for cFos-IR (red) was often detected in cells that expressed YFP (green; double arrowhead), but not in cells that were immunolabelled for PKCγ (blue). **(E)**, Many cFOS-labelled cells expressed both YFP and SOM (blue) and YFP (double arrowhead and inset), whereas others expressed only SOM (asterisk). **(F)**, Photostimulation induced cFOS expression in Pax2-expressing interneurons (asterisk; blue), with some of these cells also showing immunolabelling for YFP (double arrowhead and inset). **(G)**, Pie graph shows the proportion of photostimulation-evoked cFos expression accounted for by directly activating CR-ChR2 neurons (28.7%), and those populations recruited by CR-ChR2 activity including NK1R^+^ neurons (10.5%), SOM^+^ neurons (7.1%), and Pax2^+^ neurons (20.6%). The remaining fraction (33.1%) are likely to represent unidentified excitatory populations as they did not express Pax-2. **(H-I)**, Most excitatory synaptic inputs on to NK1R-expressing dendrites (green) in lamina I (arrows) were derived from axon terminals immunolabelled for SOM (arrowhead; blue), many of which also originated from CR-expressing cells (double arrowhead; red). Excitatory synaptic inputs on to the dendrites of lamina I NK1R-expressing cells (green) were identified using immunolabelling for Homer 1 (white; arrows). **(J-L)**, Most Homer puncta were directly apposed to axon terminals immunolabelled for VGLUT2 (arrowhead; I; red), many of which also co-expressed SOM (double arrowhead; J; blue). Scale bars (in µm): A, B = 100; C-F = 20; H and J = 10.

Two additional excitatory interneuron populations were also differentiated by protein kinase C gamma (PKCγ) and somatostatin (SOM) expression (Figure 3D-E). Immunolabelling for SOM was present in 13% of Fos^+^ cell profiles (10.7 ± 2.3 of 80.3 ± 21.2 neurons; 3 animals), however, approximately half of these also expressed YFP (5 ± 2.1 of 10.7 ± 2.3 neurons), consistent with the expected overlap between SOM and CR in lamina II neurons (Figure 3E). In contrast, we found no evidence of Fos^+^ cells that also immunolabelled for PKCγ (0 of 76.3 ± 5.8 neurons; 3 animals), implying that this population of excitatory interneurons is not postsynaptic to the ChR2-YFP cells (Figure 3D). Finally, 23% of Fos^+^ cells following spinal photostimulation were identified as inhibitory interneurons (21.3 ± 13.0 of 97 ± 28.4 neurons; 3 animals), by the expression of Pax2^+^ immunolabelling (Figure 3F). Surprisingly, however, only 2% of this inhibitory population expressed YFP (1.3 ± 1.2 of 21.3 ± 13.0; 3 animals), confirming inhibitory CR-ChR2 neurons are also recruited during spinal photostimulation but these cells are in the minority. Therefore, a large population of inhibitory interneurons is also engaged by activation of the excitatory CR^+^ population. The remaining Fos^+^ cells are most likely to be other unidentified populations of excitatory interneurons, due to their absence of Pax2 labelling, and these may include excitatory CR^+^ neurons that did not express ChR2-YFP (Figure 3G). The relative recruitment of each neurochemically defined population was also calculated yielding: 8.4% of all ChR2 neurons (21.17 ± 4.01 of 251.17 ± 43.31 neurons), 21% of all NK1R^+^ neurons (5.33 ± 2.08 of 25 ± 13.89 neurons), 5.5% of all SOM^+^ neurons (10.67 ± 2.33 of 194.67 ± 59.26 neurons), 0% of all PKCγ^+^ neurons (0 ± 0 of 159 ± 25.06 neurons), and 15.3% of all Pax2^+^ neurons (21.33 ± 13.01 of 139.33 ± 37.57 neurons). Taken together, these data show that activation of the CR-ChR2 network selectively recruits a diverse range of excitatory interneurons, inhibitory interneurons, and projection neurons.

### CR^+^/SOM^+^ neurons provide direct input to Projection neurons

Current models for dorsal horn microcircuitry place CR^+^ neurons in a polysynaptic circuit that signals through SOM^+^ neurons to activate LI projection neurons and initiate pain signalling (Peirs et al., 2015). Our data is compatible with this model as CR^+^ neuron photostimulation evoked oEPSCs in DH populations (including populations located superficial to the CR^+^ plexus), and produced robust cFos expression in both SOM^+^ neurons and putative NK1R^+^ LI projection neurons (Figure 3C). Since extensive co-localisation of CR^+^ and SOM^+^ has been reported previously(Gutierrez-Mecinas et al., 2016), it remained to be clarified how CR^+^ network activity reached projection neurons. This issue was addressed using a neuroanatomical approach with CR-Cre mice (Cr-IRES-Cre) crossed with a loxP-flanked-Synaptophysin-tdTomato reporter line (Ai34) to generate offspring where tdTomato labelled synaptic vesicles in CR^+^ neurons (CR^cre^;Ai34). This allowed us to define CR^+^ cells axon terminals with greater precision. Tissue from these animals was subsequently processed to identify putative LI PNs with NK1R^+^ labelling, excitatory synapses using immunolabelling for Homer^+^, and SOM^+^ labelling to differentiate inputs from CR^+^ only, SOM^+^ only, or CR^+/^SOM^+^ co-expressing inputs (Figure 3H-I). Using this strategy, ∼30% (31.5%; St. Dev. = ± 3.5, n=3 animals) of all Homer puncta on NK1R cells were derived from CR^+^ terminals, 4.2% (± 0.89) arising from CR^+^ only inputs, and 27.3% (± 3.63) from CR^+^/SOM^+^. Alternatively, 50.4% (± 0.82) of all homer puncta on NK1R cells associated with a SOM^+^ terminal, and of these 23.1% (± 2.93) were SOM^+^ only, and 27.3% CR^+^/SOM^+^.

SOM labelling in axon terminals is punctate and does not delineate the entire axonal bouton. To ensure that all SOM inputs on to NK1R cells were captured, we used immunolabelling for VGLUT2 to outline individual excitatory axon terminals, and determined what proportion of these express SOM in WT mice (n=3 animals). In this analysis, we found that 68.9% (± 6.63) of all Homer puncta on NK1R-expressing dendrites associate with VGLUT2 terminals (Figure 3 J-L). This indicates that the principal source of excitatory input to PNs is derived from interneurons. Most of these VGLUT2-IR boutons co-expressed SOM (78.9% ± 6.28). Similarly, of all Homer puncta on NK1R-expressing dendrites, 59.4% (± 8.39) were apposed by SOM^+^ boutons, of which most co-expressed VGLUT2 (91.7%; ± 1.96). This data shows that CR^+^ neurons provide substantial monosynaptic excitatory input to LI NK1R^+^ neurons, which largely represent PN’s, with most of these terminals also expressing SOM^+^. Thus together, CR^+^ and SOM^+^ interneurons constitute more than half of the excitatory input to LI NK1R^+^ neurons, and thus represent the principal source of excitatory input to these cells.

### CR-ChR2 neurons provide strong, direct input to Projection neurons

Given clear neuroanatomical evidence that CR^+^ neurons provided input to LI projection neurons (PNs) above, the functional impact of these connections was assessed. CR^cre^;Ai32 animals (n=2) received bilateral intracranial injections of AAV-CB7-Cl∼mCherry in the parabrachial nuclei and then following a 3-4 week incubation period spinal cord slices were prepared for targeted recordings from mCherry-labelled PNs. Under these conditions, brief full-field photostimulation (16 mW, 1 ms) applied to activate CR-ChR2 neurons produced oEPSCs in PNs that were blocked by bath applied CNQX (10 μM, n = 5). oEPSCs were observed in 65% of these recordings (13/20) indicating clear connectivity between the CR-ChR2 population and PNs (Figure 4A). These responses could be differentiated into single oEPSC events (8/13) and multiple oEPSC responses (5/13). Consistent with some PNs receiving convergent input from several CR-ChR2 neurons, multiple oEPSC responses exhibited slower rise times and half widths than single oEPSCs (rise = 9.66 ± 2.95 ms *vs.* 2.97 ± 0.45 ms, p = 0.015; halfwidth= 15.73 ± 2.16 ms *vs.* 7.76 ± 1.39 ms, p = 0.016). In contrast, both oEPSC response types occurred at similar post-photostimulation latencies (8.21 ± 2.2 ms *vs.* 7.53 ± 0.7 ms, p = 0.82) and together, these short latencies were comparable to those observed in other untargeted recordings dorsal to the CR^+^ plexus and CR-ChR2 recordings (7.95 ± 1.38 ms *vs.* 7.34 ± 1.06 ms, p = 0.981; and *vs.* 8.61 ± 1.23 ms, p = 0.985, respectively). The strength of CR-ChR2 to PN connections was also assessed in current clamp where photostimulation-mediated oEPSPs were capable of evoking an action potential in ∼70% of PNs (5/7), with a reliability of 0.675 (ie, 67.5% chance of a suprathreshold AP response). Neuroanatomical confirmation of direct CR-ChR2 derived input to PNs was also obtained in 3 of 5 PNs that were neurobiotin recovered, where clear CR-ChR2 puncta were identified in close apposition to neurobiotin labelled dendrites (Figure 4B). Together, these results demonstrate a functionally relevant monosynaptic connection exists between CR-ChR2 neurons and PNs and is capable of recruiting PN discharge.

**Figure 4.**
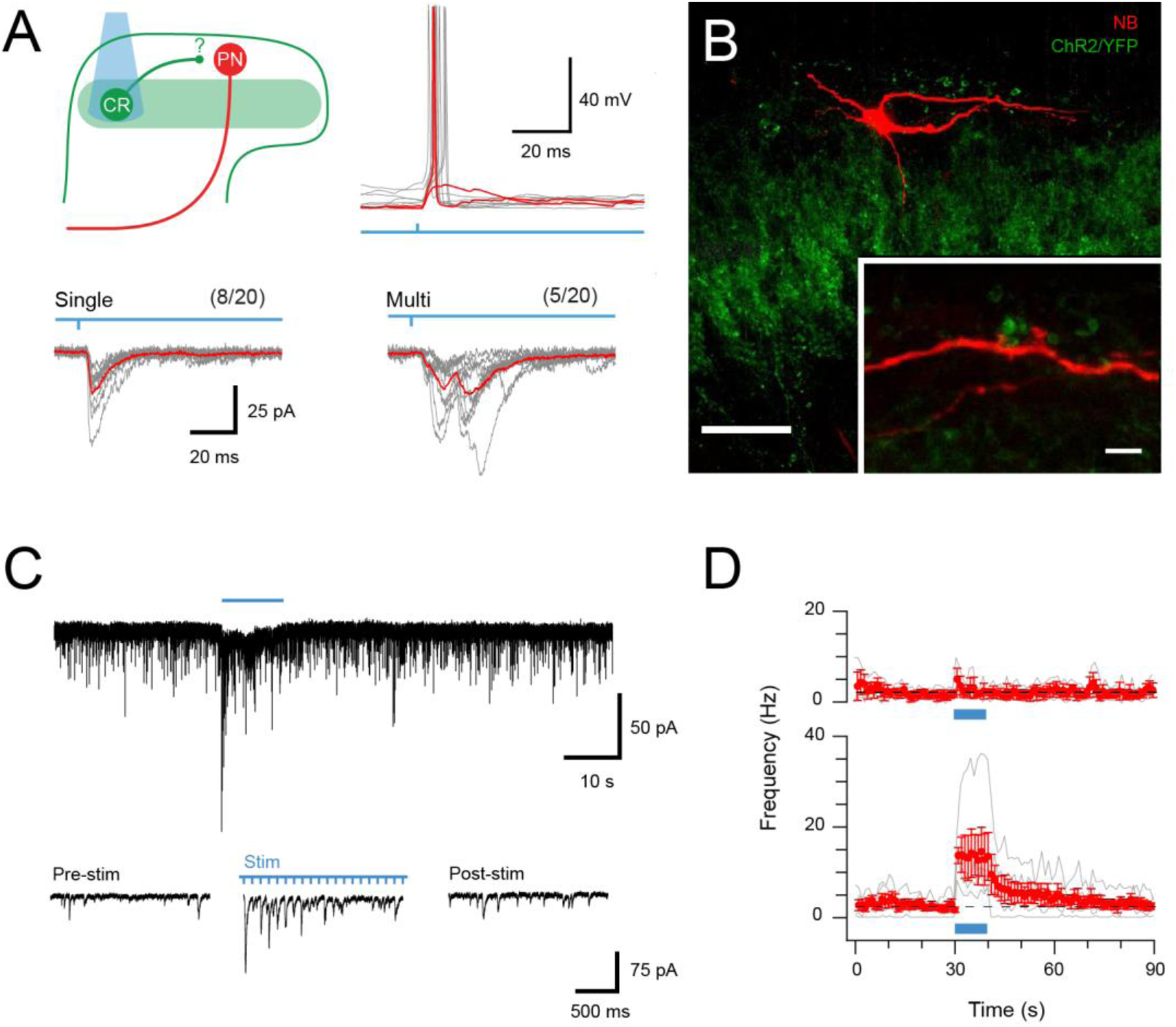
CR-ChR2 neurons provide synaptic input to LI PNs. **(A)**, Schematic (upper left) shows experimental setup with CR-ChR2 neuron (CR) photostimulation (PS) applied while monitoring retrograde virus labelled projection neuron (PN) activity (n = 20 cells from 2 animals). Lower traces show example PS evoked inward currents recorded in PNs under voltage clamp. Responses (grey) showed either single (left) or multicomponent responses (right) during PS (blue bar) with the average response overlaid (red). Values above show number of PN recordings that exhibited PS responses. Upper right traces show an example PN recording under current clamp with PS evoked input from CR neurons able to initiate AP discharge in PNs (individual subthreshold and suprathreshold responses in red). **(B)**, Image shows neurobiotin recovered PN (red) relative to expression of YFP/ChR2 in CR neurons (green). High magnification inset shows PN dendrite in close apposition with YFP/ChR2 puncta. Scale bars (in µm): 50; inset 5. **(C)**, Trace shows recording from a PN with EPSCs before, during and following full field PS of CR neurons (blue bar, 16 mW, 10 ms pulses @ 10 Hz, 10 seconds). Insets below show EPSC activity before, during and following PS on expanded time scale. Note the increase in EPSCs during and following the PS period. **(D)** Plots compare mean EPSC frequency (red) for PNs deemed to exhibit a significant PS increase (lower, n=5, EPSC frequency during PS exceeded mean baseline frequency ± 4SD), and PNs with a similar baseline EPSC frequency, but no PS evoked change in activity (upper, n =3).

In light of this connectivity, the impact of repeated CR-ChR2 photostimulation was also assessed to determine if this pathway supported the enhanced signalling seen in the CR^+^ network (see Figure 2). Under these conditions ∼60% of PNs tested (5/8) exhibited significant and sustained responses during extended CR-ChR2 photostimulation (10 s @ 10 Hz, 10 ms pulses at 16 mW), defined as an increase in 4 standard deviations above the mean background sEPSC frequency (Figure 4C). This increase reflected the stimulation features (ie, approximately 10 Hz increase) and took ∼20 s to return to baseline (Figure 4D). In contrast, the remaining PNs still exhibited CR-ChR2 input but did not show sustained responses (3/8). In conclusion, these results confirm that strong signalling arising from the CR^+^ network reaches PNs, the output cell of the DH, and therefore drive substantial output signals to higher brain regions in the ascending pain pathway.

### Photostimulation in behaving CR-ChR2 mice

The functional significance of CR-ChR2 neuron connectivity and activation within the DH was tested by chronically implanting a fiber optic probe over the surface of the dorsal spinal cord in CR^cre^;Ai32 mice for subsequent photostimulation. Behavioural responses were first tested using a range of photostimulation intensities (0.5 – 20 mW, 10 ms pulses @ 10 Hz for 10 s; Supplementary Figure 5), which produced clear behavioural responses (Supplementary Video 1). Specifically, responses were characterized by targeted nocifensive behaviour including paw lifting and licking/biting, typically focussed to the hindpaw or hindlimb region. The intensity and duration of these responses increased with photostimulation intensity until 10 mW and then stabilised above this (n=5, Supplementary Figure 5). Thus, a photostimulation intensity of 10 mW was adopted for subsequent experiments, unless otherwise noted. A larger cohort of CR^cre^;Ai32 (n=25) animals was then assessed, exhibiting nocifensive behaviour initiated at the onset of photostimulation (10 mW, 10 ms pulses @ 10 Hz for 10 s) and outlasting the photostimulation period. (Figure 5A). In contrast, photostimulation in a cohort of fiber optic probe implanted CReGFP (n=9) animals did produce photostimulation time-locked behaviours, although random grooming bouts were occasionally observed (Figure 5A).

**Figure 5.**
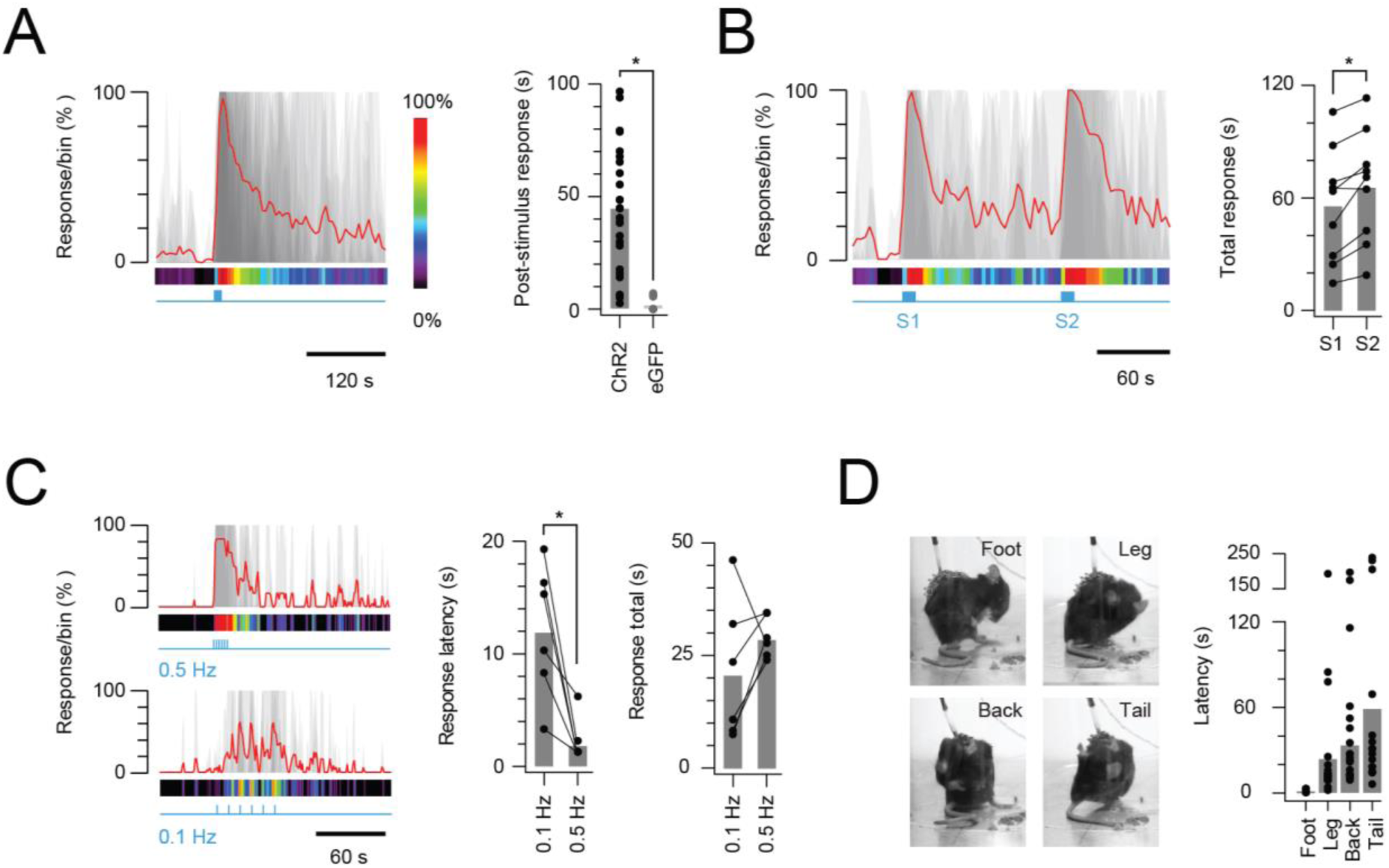
In vivo photostimulation response characteristics. **(A)**, Left plot shows overlaid peristimulus histograms of photostimulation responses in CR-ChR2 mice (n=25, grey traces) with response duration binned in 5 s epochs. Mean response is shown in red and converted to a heat bar (bins color coded to percent time groomed (red = 100%, black = 0%). Right plot compares total response duration for CR-ChR2 mice and a control group of CR-eGFP mice (n=9). Note CR-ChR2 mouse responses varied with an average of 45 seconds nocifensive behavior outlasting the 10 s photostimulation period. (**B**), Left plot shows overlaid responses from a subset of animals (n=6) that received two successive photostimuli, separated by 120 s (10 mW, 10 Hz, 10 s, grey traces). Right plot compares total response duration to initial (S1) and repeat photostimulation (S2) highlighting an average 10 s increase in the second response (p=0.005). **(C)**, Left plots show overlaid responses to brief subthreshold photostimulation trains (0.72 ± 0.36 mW, 10 ms pulses) delivered at two frequencies (0.1 Hz – lower, and 0.5 Hz – upper). Right plots compare latency to photostimulation responses and total response duration at the two stimulation frequencies. Repeated subthreshold photostimuli summate to evoke a response and latency is significantly reduced by increased stimulation frequency (p=0.012). **(D)** Photostimulation responses mapped to the body region targeted in a subset of CR-ChR2 animals (paw, leg, back or tail; n=12). Images show examples of nocifensive responses directed to the hind paw, hindlimb, back, and tail. Plots (right) compare group data for the latency of nocifensive behavior directed to different body regions. Hind paw focused responses show the shortest latency followed by significantly longer latencies for responses targeting the hind limb, back and then tail.

Given the sustained nature of post-photostimulation nocifensive responses, the potential for subsequent responses to be enhanced by prior CR-ChR2 activation was investigated by delivering two successive 10 s bouts of photostimulation, separated by 120 s (Figure 5B, n=9 animals). Under these conditions the second nocifensive response was significantly longer lasting than the first (56.26 ± 10.09 s *vs*. 66.16 ± 9.95 s p=0.005; Figure 5B). This mirrored the observation that of some CR-ChR2 neurons received sustained levels of excitatory signalling for a period following recruitment *in vitro*, and this signal reach LI PNs. To further explore summation of CR-Ch2R^+^ network activity, short trains of subthreshold photostimuli (no behavioural response to a single pulse) were delivered at two frequencies (0.1 and 0.5 Hz). Photostimulation intensities that did not evoke a behavioural response during 1 s of stimulation (10 ms pulses at 10 Hz) were first established for each animal (n=6, 0.72 ± 0.36 mW, range 0.1-2 mW). This stimulus was then repeated once every 10 s over 1 min (0.1 Hz), and once every 2 s for 12 s (0.5Hz), altering the window for summation and associated behavioural responses without changing the total energy used to activate CR-ChR2 neurons (Figure 5C). These trains of stimuli reliably evoked behavioural responses at both frequencies (0.1 and 0.5 Hz) despite the absence of responses to single stimuli. The response characteristics differed, however, in that the response latency for 0.1 Hz stimulation was significantly longer than for the higher frequency (0.5 Hz) protocol (11.83 ± 2.41 ms *vs*. 2.00 ± 0.82 ms, p=0.012). The relationship between stimulation frequency and total response was more varied with 5/6 animals exhibiting longer responses for the higher frequency stimulation (0.5 Hz) but response duration falling in one animal. Thus, response duration was statistically similar in both stimulation frequencies (20.57 ± 6.31 ms *vs.* 28.73 ± 1.86 ms, p= 0.218). This may be explained by the altered photostimulation frequency also changing the stimulation period duration and influencing the analysis. Regardless, these multiple stimulation paradigms reinforce the ability of CR^+^ neuron networks to retain subthreshold and suprathreshold excitation within an integration window that can influence the characteristics of subsequent responses.

Another pronounced feature of the CR-ChR2 photostimulation response was the dynamic nature of the area targeted for nocifensive behaviour beyond the initial (primary) body region. This observation was characterized in a subset of animals (n=18) where the initial response was directed at the right hind paw, allowing similar comparisons across multiple animals (Figure 5D). In this analysis, the onset of nocifensive responses directed to the right paw, right hind limb, back, and tail were measured, revealing a stereotypic progression of the nocifensive response across dermatomes. Comparison of the latency to nocifensive responses at each dermatome reinforced this stereotyped behaviour (right paw = 1.25 ± 0.22 s *vs*. Right hind limb = 26.42 ± 9.11 s *vs*. Back = 40.83 ± 10.78 s *vs*. Tail = 59.29 ± 18.32 s, p=0.003; Figure 5D). Thus, in addition to sustaining excitation beyond photostimulation, optogenetic activation of the CR-ChR2^+^ neuron network produced sensory signalling that spread to adjacent dermatomes before subsiding.

### CR-ChR2 photostimulation responses are nociceptive and aversive

Although CR-ChR2^+^ photostimulation responses appeared nociceptive ‘pain-like’ in quality, additional analysis was needed to support this interpretation. First, Fos-protein activity mapping detected a distinct distribution of Fos^+^ neurons in the brains of CR^cre^;Ai32 (n=5) versus CReGFP (n=5) mice following a single bout of spinal photostimulation (10 mW 10 ms pulses @ 10 Hz for 10 s). Specifically, robust Fos^+^ expression was detected in the somatosensory cortex (S1), cingulate cortex, insular cortex, and parabrachial nucleus (PBN) of CR^cre^;Ai32 mice (Figure 6A). This expression pattern was significantly elevated above the CReGFP control group (S1 p=0.043, cingulate p=0.016, insula p=0.035, and PBN p=0.023). Neuronal activity in these regions is consistent with nociceptive signalling being relayed along the neuroaxis, mirroring the pronounced nocifensive responses observed during *in vivo* photostimulation.

**Figure 6.**
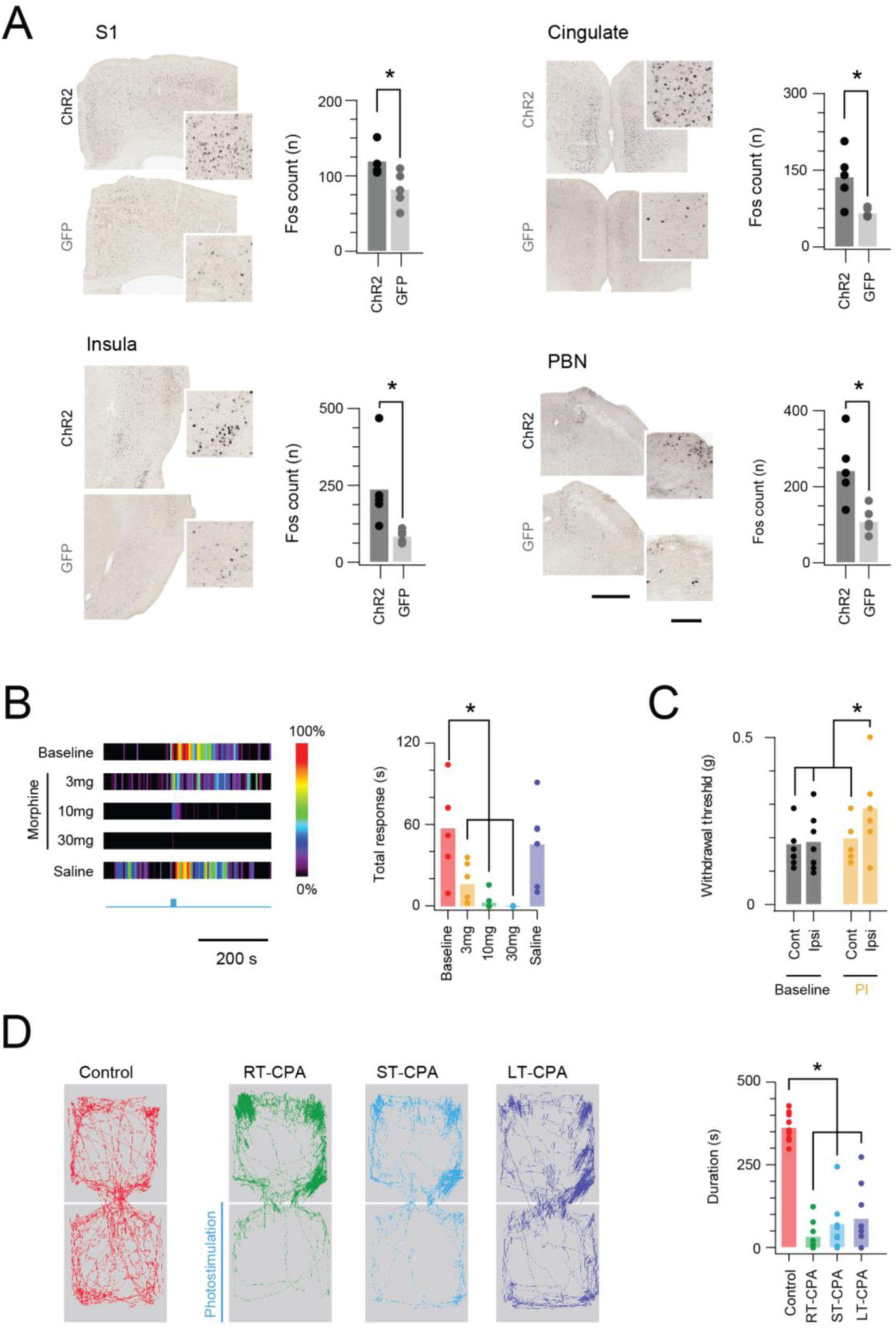
In vivo CR-ChR2 photostimulation responses have nociceptive characteristics. **(A)**, Images show representative brain sections from CR^cre^;Ai32 (upper) and CR-GFP animals (lower) immunlabelled for Fos protein following *in vivo* spinal photostimulation (10 mW at 10 Hz, 10 min), insets show Fos labelled profiles at higher magnification. Bar graphs (right) of group data compare Fos^+^ neuron counts in corresponding brain regions. The number of Fos^+^ profiles was elevated in CR^cre^;Ai32 photostimulated mice in the somatosensory cortex (S1, p=0.043), anterior cingulate cortex (Cingulate, p=0.016), Insula cortex (Insula, p=0.0346), and parabrachial nucleus (PBN, p=0.023). Scale bars = 500µm, inset = 100µm. (**B**), Heat bars (left) show average photostimulation responses (10 mW at 10 Hz, 10 s), from a subset of animals (n=6) assessed under baseline conditions, after, three morphine doses (3, 10, and 30mg/kg i.p.), and control saline injection. Bar graphs (right) compares group data for total nocifensive response duration under each condition. Morphine produced a dose dependent block of the nocifensive response compared to baseline (3mg/kg, p=0.024; 10mg/kg, p=0.002; 30mg/kg, p=0.001). (**C**), Group plots compare mean withdrawal threshold (von Frey) for hind paws ipsilateral and contralateral to spinal fiber optic probe placement. Left bars show mean withdrawal threshold under baseline conditions and right bars show mean withdrawal threshold assessed with NpHR3-mediated photoinhibition of CR^+^ neurons in the ipsilateral spinal cord. Spinal photoinhbition selectively increased withdrawal threshold for the ipsilateral hindpaw, consistent with CR^+^ neuron inhibition decreasing mechanical sensitivity (cont = contralateral, ipsi = ipsilateral). (**D**) Four maps (left) show mouse activity traces during the conditioned place aversion (CPA) testing under: control conditions (no photostimulation – red trace); real time CPA training (RT-CPA – green trace) where one arena is assigned for photostimulation (10mW, 10Hz, 10s in every minute) on entry; short-term CPA (ST-CPA), assessed 1 h following the RT-CPA session with no photostimulation (blue trace); and long-term CPA (LT-CPA), assessed 24 h after the RT-CPA session again with no photostiulation (purple trace). Bar graph (right) compares time spent in the photostimulation arena during CPA testing. Spinal photostimulation of CR-ChR2 neurons established a robust real-time conditioned place aversion by the fourth RT-CPA session (green) significantly reducing time in the photostimulation arena (p<0.0001). These reductions persisted during ST-CPA (p=0.0001) and LT-CPA (p=0.0001).

Given nocifensive responses should also be sensitive to analgesia, a group of CR^cre^;Ai32 animals (n=5) also underwent photostimulation during a randomized schedule of varying degrees of morphine analgesia (3mg/kg, 10mg/kg, or 30mg/kg morphine s.c; or vehicle only injection of saline, s.c.). Identical photostimulation was delivered under each condition (10 mW, 10 ms pulses @ 10 Hz for 10 s) and behavioural responses analysed to provide a robust assessment of analgesic sensitivity (Figure 6B). Morphine produced a dose dependent reduction in the photostimulation-induced nocifensive behaviour, which was abolished at the highest morphine dose (30mg/kg). This reinforces the nocifensive nature of the circuits activated during CR-ChR2^+^ photostimulation.

While the above data shows CR-ChR2^+^ photostimulation was sufficient to evoke responses consistent with nociceptive pain, this did not confirm necessity of the CR^+^ network in sensory-evoked responses. Thus, a photoinhibition approach was also employed crossing CR-cre mice with loxP-flanked-NpHR3eYFP mice (Ai39) to generate offspring where halorhodopsin (NpHR3) was expressed in CR^+^ neurons (CR^cre^;Ai39). To validate the expression of NpHR3-YFP expression in CR cells, we assessed the incidence of co-expression of these markers in spinal neurons in laminae I and II (n = 3 animals). We found that 82.2% (± 1.27) of YFP-expressing cells were immunopositive for CR (967 YFP cells analysed; range 281, 316 and 370 cells per animal; Supplementary Figure 6A), and that 94.1% (± 4.27) of CR-IR cells expressed YFP (225 CR-IR cells analysed; 58, 78 and 89 cells per animal). Furthermore, full field illumination (590 nM, 20 mW) evoked prominent outward currents consistent with a NpHR3-mediated potassium conductance (Supplementary Figure 6B). NpHR3-mediated photoinhibition of AP discharge was confirmed by comparing CR-NpHR3^+^ neuron spiking responses to depolarizing current injection with and without photoinhibition (Supplementary Figure 6C). Under these conditions photoinhibition increased the rheobase current required to activate AP spiking and decreased the number of APs evoked during increasing current steps (Supplementary Figure 6C). CR;Ai39 mice (n=8) were subsequently implanted with spinal fiber optic probes and underwent von Frey mechanical threshold testing with and without NpHR3-mediated photoinhibition (590 nM, 20 mW) in alternating order over 4 days (Figure 6C). *In vivo* photoinhibition significantly reduced paw withdrawal threshold on the ipsilateral (photoinhibited) hind paw (0.19 ± 0.02 g *vs.* 0.29 ± 0.04 g, p=0.017) but not contralateral side (0.18 ± 0.02 g *vs.* 0.19 ± 0.03 g, p=0.721), confirming a role for the CR^+^ network in setting withdrawal thresholds (Figure 6C).

Finally, the relative potency and valence of spinal photostimulation was assessed in a group of CR^cre^;Ai32 animals (n=13) with spinal fiber optic probes implanted that subsequently underwent conditioned place aversion testing (Figure 6D). Baseline preference for each animal was determined in a two-arena enclosure without photostimulation, and the preferred arena was then assigned for photostimulation (10 mW 10 ms pulses @ 10 Hz for 10 s in every min). Animals were subsequently tested for a real-time place aversion (RT-PA) over 4 sessions, i.e. learned avoidance of the photostimulation arena (Supplementary Video 2). Animals that exhibited a strong RT-PA on the last two sessions, defined as a 50% reduction from baseline in time spent in photostimulation area, were subsequently tested for a traditional conditioned place aversion (CPA) 1 h (short term) and 24 h (long term) after the last RT-PA session (9/13 animals). Importantly, no photostimulation was delivered during this CPA testing, instead assessing the aversive nature of photostimulation recall. In short term CPA testing (ST-CPA), animals retained a significant aversion to the previous photostimulation arena compared to baseline (363 ± 15 s *vs*. 72 ± 25 s, p=0.0001). Likewise, in long term CPA testing (LT-CPA) aversion to the previous photostimulation arena was still apparent 24 h after RT-PA (363 ±15 s *vs.* 89 ± 31 s, p=0.0001). Thus, spinal photostimulation of the CR-ChR2^+^ population produced a potent sensory experience with a strong and lasting negative valence.

## Discussion

Our limited understanding of sensory coding in the spinal cord remains a significant barrier to defining how normal sensory experience evolves and how pathological conditions such as chronic pain develop(Todd, 2010, Hachisuka et al., 2018). In this study we applied both *in vitro* and *in vivo* optogenetic approaches and show that a specific population of DH interneurons that express CR form a highly interconnected excitatory network that is capable of driving excitation in multiple postsynaptic DH neuron populations, including direct connections to projection neurons. Excitation of this microcircuitry outlasts initial CR^+^ neuron activation, indicating the CR^+^ network of reciprocal excitatory connections has the capacity to sustain synaptic activity. Extending these *in vitro* findings, optogenetic activation of the CR^+^ population in awake animals caused a profound and multifaceted nocifensive response, indicating that this network can prolong, spread and amplify spinal nociceptive signaling. Together, these findings provide a detailed examination of the CR^+^ excitatory interneuron population and the postsynaptic circuits they activate during sensory processing.

### CR^+^ neuron microcircuits

Our *in vitro* electrophysiology showed that CR^+^ neurons exhibit diverse functional excitatory synaptic connections within the DH. A surprising finding in this work was the high degree of interconnectivity between CR^+^ neurons, establishing an excitatory network that when recruited, could substantially enhance excitation in the DH (Figure 1 and 2). This is in line with observations using viral-mediated excitatory DREADD-expression to activate CR^+^ neurons selectively (Peirs et al., 2015). This work showed that CNO exposure activated a substantial proportion of transduced DREADD-positive CR^+^ neurons (30%), however, an additional large proportion of DREADD-negative CR^+^ neurons (45%) were also activated. This supports the capacity of CR^+^ neuron recruitment to drive activation of the wider CR^+^ network. Such interconnectivity in a neurochemically defined population has been described for another DH population identified by expression of neurotensin (Hachisuka et al., 2018), although this work did not go on to demonstrate the functional relevance of this observation in behaving animals. Nevertheless, this work highlighted how interconnected excitatory networks could potentiate excitatory DH signaling. Likewise, similar arrangements of interconnected neuronal networks have been reported in other CNS regions, albeit typically involving inhibitory populations (Tamas et al., 2000, Woodruff and Sah, 2007, Meyer et al., 2002). This interconnectivity among excitatory interneurons suggests the CR^+^ network represents a potent source of excitatory signaling, that could amplify input in the DH.

CRACM experiments also showed that input from CR^+^ neurons is widespread, with other neurons located in the CR^+^ plexus as well as more dorsal populations, receiving short and longer latency input following CR-ChR2 activation. These experiments also confirmed that one of the dorsal populations to receive this input was LI projection neurons, which often received convergent, multicomponent inputs and more reliably discharged APs in response to this input than other populations. A slower time course in these projection neuron responses also suggests summation of several asynchronous inputs through multiple pathways. These features are consistent with CR^+^ neurons initiating excitation that converges on projection neurons. Such excitatory relays have been proposed in the DH as a substrate for allodynia, where low threshold mechanical inputs are transmitted into nociceptive circuits (Peirs et al., 2015, Torsney and MacDermott, 2006, Miraucourt et al., 2007, Neumann et al., 2008), as well as feed forward circuits that supplement excitation during nociceptive processing (Lu et al., 2013, Lu and Perl, 2005). For example, Peirs et al., (2015) showed that CR^+^ neurons receive low-threshold input via a population of lamina III neurons that transiently express VGLUT3. They also used Fos labelling and DREADD silencing of various populations to propose a circuit connecting CR^+^ neurons to projection neurons in lamina I via polysynaptic connections that included an interposed population of SOM^+^ interneurons. In support of this proposal, we find approximately 30% of all excitatory input to lamina I projection neurons are derived from axons of CR^+^ interneurons (Figure 3), of which a significant proportion also express SOM (86.5%; ± 3.17). These findings demonstrate that CR^+^ neurons provide considerable direct input to the lamina I projection neuron population, and when considered alongside the CR^+^ network interconnectivity, these have the potential to greatly influence spinal sensory outputs. Distinctions in these predicted circuits (monosynaptic from CR^+^ neurons versus polysynaptic via SOM^+^ neurons) may reflect methodological differences, with the Peirs work using a viral strategy that identified a relatively narrow population of CR^+^ neurons, whereas our experiments used a transgenic breeding approach identifying a much larger CR^+^ population. In agreement with Peirs et al (2015), however, our dataset suggests excitatory SOM^+^ and projection neurons are postsynaptic to the CR^+^ population, but not PKCγ^+^ excitatory interneurons (Figure 3). Importantly, the high fidelity of YFP expression with CR immunolabelling in our mouse lines validates that our behavioral observations and circuit diagrams are the result of manipulating CR-expressing cells, and not contaminated by populations of unidentified neurons that express CR^+^ transiently during earlier developmental time-points. In summary, the interconnectivity of CR^+^ neurons, combined with the postsynaptic circuitry we have identified provides a mechanism to relay low threshold input into nociceptive circuits (allodynia), and provide additional excitation during nociceptive processing (hyperalgesia) through reverberating patterns of excitation.

Inhibitory signaling has been a central element in models of spinal sensory processing since publication of the gate control theory and contemporary work from a number of groups has since identified several critical inhibitory populations (Petitjean et al., 2015, Duan et al., 2014, Foster et al., 2015, Cui et al., 2016). Consistent with these views, our Fos mapping also supports a role for the CR^+^ network in engaging inhibitory interneurons. These inhibitory circuits may be important for modality coding by suppressing selective populations while the CR^+^ circuits are active (Zeilhofer et al., 2012, Price and Prescott, 2015). Such inhibition is known to act in other sensory systems to refine receptive field characteristics and a similar constraint over the activation of CR^+^ neurons would fit such a model (Woolf and Fitzgerald, 1983, Kato et al., 2011). Finally, ongoing inhibition can tune sensory thresholds, most evident in the wealth of data showing that diminished inhibition leads to pathological conditions such as chronic pain and itch (Moore et al., 2002, Coull et al., 2003, Zeilhofer, 2005, Ross et al., 2010). An important distinction between the current study and previous work is that the inhibition evoked in our study is driven exclusively by recruitment of CR^+^ spinal interneurons, rather than as a result of primary afferent drive. Thus, the CR^+^ related microcircuits appear to have an inbuilt mechanism to limit the outcome of their activity under control conditions. By extension, any reduction to this inhibition would unmask added excitation with DH circuits with relevance to many pathological conditions that feature aberrant excitation. We show that CR^+^ network evoked inhibition is widespread in the DH, with a range of latencies that indicate both direct and indirect circuits are engaged (Supplementary Figure 4). Direct inhibition is not surprising given we have previously described a small inhibitory CR^+^ population (Smith et al., 2015), and show here that they express ChR2 in the CR^cre^;Ai32 animals used. These neurons are directly activated during spinal photostimulation and therefore provide the only source of short latency monosynaptic inhibition. In contrast, secondary recruitment of other inhibitory populations by the excitatory CR^+^ neurons produces longer latency polysynaptic inhibition. Previous work has shown dynophin^+^ neurons provide important gating inhibition to somatostatin^+^ neurons(Duan et al., 2014), and given the overlap between somatostatin and CR, CR/SOM cells are therefore also likely to contribute to the polysynaptic inhibition observed here. Similarly, GABAergic enkephalin^+^ neurons have recently been implicated in gating mechanical pain (Francois et al., 2017) and thus may also contribute to CR^+^ network evoked inhibition.

### Behavioral consequence of CR-ChR2 activation

The behavioral consequence of experimentally activating CR^+^ circuits *in vivo* was striking, with a targeted and multifaceted nocifensive response (paw licking, biting, shaking) directed to dermatomes predicted by the location of fiber optic implant and initiated upon photostimulation (Supplementary Figure 5). This resembles reports from two other groups that have applied spinal photostimulation to populations of DH neurons. The most relevant assessed excitatory somatostatin^+^ neurons, reporting photostimulation produced abrupt nocifensive behavior and a conditioned place aversion (Christensen et al., 2016). This conserved behavioral profile is again consistent with the overlap in somatostatin and CR populations studied and the postsynaptic position of somatostatin^+^ neurons relative to the CR^+^ population. The Christensen et al (2016) study, however, also unmasked an itch related behavior using modified photostimulation parameters. We confirmed the nociceptive nature of CR^+^ photostimulation by demonstrating selective Fos labelling in distinct pain processing brain nuclei as well as sensitivity of the behavioral responses to morphine (Figure 6). We observed a dose dependent inhibition of photostimulation related behavior that was abolished at the highest morphine dose. Importantly, morphine is a prototypical analgesic but does not have antipruritic actions, in fact it induces scratching when administered intrathecally, independent of analgesia (Lui and Ng, 2011). Thus, coupled with the robust neuronal activation we report in key regions in the ascending pain pathway (Figure 6), this work reinforces CR^+^ activated pathways are nociceptive in quality.

It is worth noting that a small number of inhibitory CR^+^ neurons were also activated during photostimulation (Figure 3G). Previous optogenetic manipulation of inhibitory populations in the DH has only been reported using archaerhodopsin, which produced a predictable decrease in sensory thresholds(Bonin et al., 2016). The related approach of chemogenetic activation has, however, been applied to activate parvalbumin^+^ inhibitory interneurons in the DH, demonstrating an increase in sensory thresholds as well as an attenuation of nerve injury induced allodynia (Petitjean et al., 2015). Taken together, these results support the well-established role of inhibitory populations in suppressing spinal nociceptive signaling and suggest the recruitment of some inhibitory CR^+^ neurons in our experiments will have had minimal effects on the robust behavioral outcomes attributed to the excitatory CR^+^ population. Furthermore, we show that halorhodopsin-mediated inhibition or the CR^+^ population increased mechanical withdrawal thresholds, consistent with the result that would be predicted for inactivation of an inhibitory population from the above work.

Three behavioral observations during CR^+^ neuron photostimulation warrant further discussion. First, behavioral responses persisted well beyond the termination of photostimulation (Figure 5, Supplementary Figure 5). Coupled with the interconnectivity of CR^+^ neurons and the sustained increased in spontaneous excitatory activity under extended *in vitro* photostimulation conditions, this indicates signaling can be maintained by this excitatory network. Second, behavioral responses to repeated photostimulation were enhanced, indicating short-term plasticity within the CR^+^ network (Figure 5). Previous work has focused largely on plasticity between primary afferents and DH populations(Baba et al., 2001, Luo et al., 2014), whereas our findings extends our understanding to show plasticity also occurs within intrinsic DH circuits. Finally, photostimulation responses showed a predictable pattern that initially saw nocifensive behavior focused on a specific body region (commonly the paw) but then progressed over a number of adjacent dermatomes (Figure 5). This, coupled with spinal Fos activation patterns, which extended over the mediolateral extent of the DH (Figure 3), suggest the excitatory CR^+^ network provides a pathway to spread excitation across normal dermatome boundaries. Together, these observations extend on the current view of excitatory interneurons in spinal sensory processing, which ascribes a relative limited role of linking low threshold modality (innocuous) tactile input to excite more dorsal nociceptive circuitry (Peirs and Seal, 2016, Duan et al., 2014, Yu et al., 2017). We suggest these neurons also provide a polysynaptic network to enhance/amplify local excitation, prime the region for subsequent responses, and spread excitatory signaling across modality borders.

### Conclusions

The results from this study confirm that CR^+^ neurons form an excitatory DH network that contributes to spinal pain signaling and can amplify pain signals in the absence of peripheral input. The strong interconnectivity of this neurochemially-defined subpopulation in the DH means they are ideally positioned to alter incoming sensory information prior to its relay to higher brain centers. This capacity is clearly demonstrated in the nocifensive responses elicited by spinal photostimulation. Importantly, however, our work has largely focused on characterizing CR^+^ microcircuits and establishing their functional roles in sensory experience by experimental activation using optogenetics. Future work must continue to determine how these circuits respond during peripherally evoked sensory processing, as we have here using halorhodopsin. Furthermore, the question of how pathology and injury can recruit or alter CR^+^ circuits will also be critical for determining how they contribute to symptoms of pathological pain, and how best to target them for therapeutic benefit.

## Experimental Procedures

### Animals and Ethics

Optogenetic studies were carried out on mice derived by crossing Calb2-IRES-cre (Jackson Laboratories, Bar Harbor, USA; #010774) with either Ai32 (Jackson Laboratories, Bar Harbor, USA; #024109) or Ai39 (Jackson Laboratories, Bar Harbor, USA, #014539) to generate offspring where ChR2/YFP or NpHR/YFP was expressed in CR^+^ cells (CR^cre^;Ai32 or CR^cre^;Ai39). Axon terminal labelling experiments crossed Calb2-IRES-cre and Ai34D (Jackson Laboratories, Bar Harbour, USA, # 012570) mice to generate offspring with CR^+^ axon terminals labelled with TdTomato. In control experiments, another transgenic mouse line with enhanced green fluorescent protein expressed under the control of the calretinin promoter (CReGFP) was used (Caputi et al., 2009). All experimental procedures were performed in accordance with the University of Newcastle’s animal care and ethics committee (protocols A-2013-312 and A-2016-603). Animals of both sexes were used for electrophysiology (age: 3-12 months) and behavior experiments (age: 8-12 weeks).

### Spinal slice preparation

Acute spinal cord slices were prepared using previously described methods (Graham et al., 2003, Graham et al., 2011). Briefly, animals were anaesthetized with ketamine (100mg/kg i.p) and decapitated. The ventral surface of the vertebral column was exposed and the spinal cord rapidly dissected in ice-cold sucrose substituted cerebrospinal fluid (ACSF) containing (in mM): 250 sucrose, 25 NaHCO_3_, 10 glucose, 2.5 KCl, 1 NaH_2_PO_4_, 1 MgCl and 2.5 CaCl_2_. Either parasagittal or transverse slices were prepared (L1-L5, 200µm thick: LI-L5 300μm thick, respectively) both using a vibrating microtome (Campden Instruments 7000 smz, Loughborough, UK). Targeted CR^+^ and unidentified recordings were undertaken in parasagittal slices, whereas targeted PN recordings used slices in the transverse plane. Slices were transferred to an interface incubation chamber containing oxygenated ACSF (118mM NaCl substituted for sucrose) and allowed to equilibrate at room temperature for at least one hour prior to recording.

### Patch clamp electrophysiology

Following incubation, slices were transferred to a recording chamber and continuously superfused with ACSF bubbled with carbanox (95% O_2_, 5% CO_2_) to achieve a final pH of 7.3-7.4. All recordings were made at room temperature. Neurons were visualised using a 40x objective and near-IR differential interference contrast optics. To identify CR-ChR2/CR-NpHR^+^ neurons, which expressed YFP, slices were viewed under fluorescence using a FITC filter set (488nm excitation and 508nm emission). CR^+^ neurons were concentrated within LII of the DH, described previously (Smith et al., 2015, Smith et al., 2016), and is easily identified as a plexus of YFP fibers and soma under fluorescent microscopy. All recordings were made either within or dorsal to this CR^+^ plexus. The parasagittal slicing approach allowed easy differentiation of the two CR populations we have previously described (Smith et al., 2015, Smith et al., 2016). Specifically, the CR^+^ excitatory population exhibits a restricted dendritic profile, whereas less common inhibitory CR^+^ neurons possess extensive rostro-caudal projecting dendritic arbours. Patch pipettes (4-8 MΩ) were filled with either a potassium gluconate based internal for recordings of excitatory input and action potential (AP) discharge, containing (in mM): 135 C_6_H_11_KO_7_, 6 NaCl, 2 MgCl_2_, 10 HEPES, 0.1 EGTA, 2 MgATP, 0.3 NaGTP, pH 7.3 (with KOH); or a caesium chloride-based internal solution for inhibitory input recordings, containing (in mM): 130 CsCl, 10 HEPES, 10 EGTA, 1 MgCl_2_, 2 MgATP and 0.3 NaGTP, pH 7.35 (with CsOH). Neurobiotin (0.2%) was included in all internal solutions for *post-hoc* cell morphology. All data were acquired using a Multiclamp 700B amplifier (Molecular Devices, Sunnyvale, CA, USA), digitized online (sampled at 10-20 kHz, filtered at 5-10 kHz) using an ITC-18 computer interface (Instrutech, Long Island, NY, USA) and stored using Axograph X software (Molecular Devices, Sunnyvale, CA, USA).

AP discharge patterns were assessed in current clamp from a membrane potential of ∼ −60mV by delivering a series of depolarising current steps (1 s duration, 20 pA increments). AP discharge was classified using previously described criteria (Graham et al., 2004, Graham et al., 2007). Briefly, delayed firing (DF) neurons exhibited a clear interval between current injection and the onset of the first AP; tonic firing (TF) neurons exhibited continuous repetitive AP discharge for the duration of the current injection; initial bursting (IB) neurons were characterised by a burst of AP discharge at the onset of the current injection; and single spiking (SS) neurons only fired a single AP at the beginning of the current step. Input resistance and series resistance were monitored throughout all recordings and excluded if either of these values changed by more than 10%. No adjustments were made for liquid junction potential. The subthreshold currents underlying AP discharge were assessed using a voltage-clamp protocol that delivered a hyperpolarizing step to −100 mV (1 s duration) followed by a depolarizing step to −40 mV (200 ms duration) from a holding potential of −70 mV. This protocol identifies four major ionic currents previously described in DH neurons, including the outward potassium currents (rapid and slow I_A_) and the inward currents, T-type calcium and non-specific cationic current I_h_.

### In vitro optogenetics

Photostimulation was achieved using a high intensity LED light source (CoolLED pE-2, Andover, UK) delivered through the microscopes optical path and controlled by Axograph X software. Recordings from excitatory versus inhibitory CR-ChR2 neurons were distinguished using their morphology in the parasagittal slice and distinct electrophysiological profiles(Smith et al., 2015, Smith et al., 2016). Photocurrents were first characterised in CR-ChR2 neurons using a current versus light intensity (488 nm, 1 second duration) analysis in voltage clamp mode. Combinations of neutral density filters were used to reduce photostimulation intensity (0.039 – 16 mW). To assess the ability and reliability of photostimulation to evoke AP discharge in CR-ChR2^+^ neurons the recording mode was switched to current clamp and brief photostimuli (16 mW, 1 ms) were delivered at multiple frequencies 5Hz, 10Hz and 20Hz (1 s duration). We then used channelrhodopsin-2 assisted circuit mapping (CRACM) to characterise the connectivity of CR^+^ neurons within the DH. The postsynaptic circuits receiving input from CR-ChR2^+^ neurons were characterized by delivering photostimulation (16 mW, 1 ms) every 12 seconds during patch clamp recordings and assessing current responses for photostimulation associated synaptic input. These recordings were made from CR-ChR2^+^ neurons as well as 3 populations of CR-ChR2 negative neurons (i.e. those that did not exhibit YFP expression). These DH neuron populations were classified relative to the distinct CR^+^ plexus within LII as either: 1) Plexus – within the CR^+^ plexus; 2) Dorsal – dorsal to CR^+^ plexus; or 3) Projection neurons – dorsal to the CR^+^ plexus and retrograde labelled (see below). The response of these populations was also assessed during prolonged photostimulation using two stimulus paradigms: 1) a 2 second continuous photostimulation (16 mW); and 2) a 1 second photostimulation (16 mW, 1 ms pulses @ 10 Hz for 1 s).

### Projection neuron recordings

To identify Lamina I projection neurons in slice recording experiments a subset of animals (n=2) underwent surgery to inject a viral tracer, specifically AAV-CB7-Cl∼mCherry, into the parabrachial nucleus (PBN). Retrograde transport of virus particles, incorporation into the genome and the subsequent expression of the mCherry protein within the PNs allowed this population to be targeted for patch clamp recording. Briefly, mice were anaesthetised with isoflurane (5% induction, 1.5-2% maintenance) and secured in a stereotaxic frame (Harvard Apparatus, Massachusetts, U.S.A). A small craniotomy was performed and up to 700nL of the viral sample was injected using a picospritzer (PV820, WPI, Florida, USA) into the PBN bilaterally. These injections were made 5.25mm posterior to bregma, ± 1.2mm of midline and 3.8mm deep from skull surface, using coordinates refined from those in the mouse brain atlas (Paxinos and Franklin, 2001). Injections were made over 5 minutes and the pipette left in place for a further 7-10 minutes to avoid drawing the virus sample along the pipette track. Animals were allowed to recover for 3 weeks to allow sufficient retrograde labelling of projection neurons before spinal cord slices were prepared. CRACM was then performed as above for other DH populations. The brain from each animal was also isolated and brainstem slices containing the PBN were prepared to confirm the injection site, which was appropriately focussed on PBN in all cases. Spinal cord slices were obtained using methods described above (*spinal slice preparation*) and mCherry positive neurons were visualised for recording using a Texas Red filter set (549 excitation, 565 emission).

### Patch clamp data analysis

All electrophysiology data were analysed offline using Axograph X software. ChR2 photocurrent amplitudes were measured as the difference between baseline and the steady state portion of the photocurrent. Excitatory and inhibitory photostimulation-evoked synaptic currents elicited by brief photostimulation, hereafter termed optical postsynaptic currents (oEPSCs and oIPSCs), were captured episodically and averaged (10 trials). Peak amplitude, rise time (10-90% of peak) and decay time constant (10-90% of the decay phase) were measured from average oEPSCs and oIPSCs. Response latency was also measured on averaged records, as the time between the onset of photostimulation and onset of the oEPSC/oIPSC. In photostimulation responses that contained multiple components a semi-automated peak detection procedure was used to determine the latency of all responses. To differentiate direct (monosynaptic) input from indirect (polysynaptic) response components the photostimulation recruitment time for CR-ChR2^+^ neurons was determined as the latency between the onset of photostimulation and the onset of AP discharge. In addition, the average time between spiking in a presynaptic neurons and a monosynaptic response in synaptically connected neurons was taken from previous paired recording studies in the spinal DH (Santos et al., 2007, Lu and Perl, 2003, Lu and Perl, 2005). These data account for the combination of AP conduction and synaptic delay that takes ∼2.5ms. Thus, windows were set for oEPSCs and oIPSCs to be considered monosynaptic by adding the photostimulation recruitment time, conduction and synaptic delays (± 2 standard deviations) of photostimulation recruitment time. Responses outside these windows were considered to more likely arise from polysynaptic activity. For longer photostimulation paradigms both oEPSCs and spontaneous excitatory postsynaptic currents (sEPSCs) were detected using a sliding template method (a semi-automated procedure in the Axograph package). Average oEPSC/sEPSCs frequency was calculated over 100ms epochs by multiplying the number of events in each epoch by 10.

To isolate oEPSCs and oIPSCs in CR-ChR2^+^ neurons, the photocurrents were first subtracted using a pharmacological approach. For oEPSCs (K^+^ gluconate-based internal), photocurrents were isolated following application of CNQX (10 μM) and then scaled to the peak photocurrent before drug application. The isolated photocurrent was then subtracted from the pre-CNQX traces leaving the isolated synaptic response (Supplementary Figure 3). The same procedure was repeated for oIPSCs (CsCl-based internal solution), except responses were obtained under 3 conditions following sequential application of CNQX (10 μM), bicuculline (10 μM), and strychnine (1 μM). In this case, the isolated photocurrent (recorded in CNQX, bicuculline and strychnine) was subtracted from the photostimulation responses under each drug condition.

### Optogenetic stimulation for Fos activation mapping

The postsynaptic circuits targeted by CR^+^ neurons were assessed by delivering spinal photostimulation to anaesthetised CR^cre^;Ai32 animals (and CReGFP control animals) and then processing spinal cords for Fos-protein and a range of additional neurochemical markers. Animals (n=5) were anaesthetised with isoflurane (5% initial, 1.5-2% maintenance) and secured in a stereotaxic frame. A longitudinal incision was made over the T10-L1 vertebrae and a laminectomy was performed on the T13 vertebra. Unilateral photostimulation (10 mW, 10 ms pulses @ 10 Hz for 10 min) was then delivered to the exposed spinal cord by positioning an optic fiber probe (400 nm core, 1 mm fiber length, Thor Labs, New Jersey, U.S.A) above the spinal cord surface using the stereotaxic frame. Photostimulation was delivered by a high intensity LED light source attached to the probe via a patch cord. Following photostimulation animals remained under anaesthesia for a further 2 hrs for subsequent comparison of Fos expression in neurochemically defined DH neurons. Animals were then anaesthetised with ketamine (100 mg/kg i.p) and perfused transcardially with saline followed by 4% depolymerised formaldehyde in 0.1M phosphate buffer. Sections were processed for immunocytochemistry by incubating in a cocktail of antibodies including chicken anti-GFP and goat anti-cFos, with either rabbit anti-NK1R, rabbit anti-somatostatin, rabbit anti-PKCγ or rabbit anti-Pax2. Full details of primary antibodies are provided in Table 1. Primary antibody labelling was detected using species-specific secondary antibodies conjugated to rhodamine, Alexa 488, Alexa 647 (Jackson Immunoresearch, West Grove, PA, USA). or with NK1-immunolabelling was visualised using a biotinylated anti-rabbit antibody (Jackson Immunoresearch) followed by a Tyramide signal amplification step using a tetramethylrhodamine kit (PerkinElmer Life Sciences, Boston, MA, USA), as described previously(Hughes et al., 2013).

### Transgenic axon terminal labelling

Analysis of CR^+^ neuron input to putative projection neurons was undertaken in tissue from CR^cre^;Ai34 mice that selectively labelled CR^+^ axon terminals with tdTomato. Animals were anaesthetised with sodium pentobarbitone (30 mg/kg *i.p.*) and perfused transcardially with Ringer solution followed by 4% depolymerised formaldehyde in 0.1M phosphate buffer. Sections were processed for immunohistochemistry by incubating in cocktails of antibodies including chicken anti-GFP, goat anti-calretinin, goat anti-Homer1, rat anti-mCherry, rabbit anti-NK1R, guinea pig anti-somatostatin, and mouse anti-VGLUT2. For full details of these antibodies, see Table 2. Primary antibody labelling was detected using species-specific secondary antibodies conjugated to rhodamine, Alexa 488, Alexa 647 (Jackson Immunoresearch, West Grove, PA, USA).

**Table 2:** Primary Antibody Details.

| Antibody | Species | Epitope | Cat no | RRID | Manufacturer | Ref | Dilution |
| --- | --- | --- | --- | --- | --- | --- | --- |
| Calretinin | Goat | Human recombinant calretinin | CG1 | AB_10000342 | SWANT | (Schiffmann et al., 1999) | 1:1000 |
| cFOS | Goat | Recombinant full-length protein | sc-52-G |  | Santa Cruz Biotechnology | (Ganley et al., 2015) | 1:2000 |
| GFP | Chicken | Recombinant full-length protein | Ab13970 |  | Abcam | (Ganley et al., 2015) | 1:1000 |
| Homer1 | Goat | Amino acids 1–175 of mouse Homer 1 | Homer1-Go-Af1270 | AB_2631104 | Frontier Science | (Nakamura et al., 2004) | 1:1000 |
| mCherry | Rat | Full-length protein mCherry | M11217 | AB_2536611 | Thermo Fisher Scientific | (Schwarz et al., 2015) | 1:1000 |
| NK1R | Rabbit | C-terminus of NK1R of rat origin, amino acids 393-407 | S8305 |  | Sigma-Aldrich | (Ptak et al., 2002) | 1:1000 |
| Pax2 | Rabbit | Amino acids 188-385 of the mouse protein | 716000 |  | Life Technologies | (Gutierrez-Mecinas et al., 2017) | 1:1000 |
| PKCγ | Rabbit | C-terminus of the mouse protein | sc211 | | Santa Cruz Biotechnology | (Gutierrez-Mecinas et al., 2016) | 1:1000 |
| Somatostatin | Guinea pig | Somatostatin-24 | IS-3/51 |  | P. Ciofi | (Ciofi et al., 2006) | 1:1000 |
| Somatostatin | Rabbit | Somatostatin-28 and somatostatin-25 | T-4103 |  | Peninsula | (Proudlock et al., 1993) | 1:1000 |
| VGLUT2 | Mouse | C-terminal sequence of rat VGLUT-2 | MB5504 | AB_2187552 | Millipore | (Hrabovszky et al., 2006) | 1:500 |

### Calretinin inputs onto filled Projection Neurons

As noted above, 0.2% neurobiotin was included in all internal recording solutions to recover recorded cell morphology. In the AAV-mediated targeted recordings of LI projection neurons, the cells were recovered using a streptavidin∼Cy5 secondary antibody before initial imaging (z=1μm, scan speed 400, pinhole 1AU) using a water immersion 25x objective on a Leica TCS SP8 scanning confocal microscope equipped with Argon (458, 488, 514nm), DPSS (561nm) and HeNe (633) lasers. Slices that contained recovered PNs were reacted with chicken anti-GFP (see Table 2 for details), to resolve axon boutons of CR neurons in close apposition with labelled PNs. Spinal slices were re-sectioned to 50μm thickness, mounted in glycerol and imaged using both a 40x oil and 63x water immersion objective. Boutons were identified as rounded YFP-labelled profiles directly apposed to labelled PN dendrites.

### Optogenetic probe surgery

Animals were anaesthetised with isoflurane (5% initial, 1.5-2% maintenance) shaved over the thoracolumbar vertebral column, secured in a stereotaxic frame and the surgical site was cleaned with chlorhexadine. Using aseptic procedures, a 3 cm incision was made over the T10-L1 vertebrae and paraspinal musculature removed. The intervertebral space between T12 and T13 was cleared to expose the spinal cord and overlying dura. Surgical staples were attached to the corresponding T12 and T13 to provide a rigid fixation point of attachment for the fiber optic probe. A probe (400 nm core, 1 mm fiber length, Thor Labs, New Jersey, U.S.A) was then positioned over the exposed spinal cord, between the staples, and fixed in place using orthodontic crown and bridge cement (Densply, Woodbridge, Canada). The surgical site was closed with sutures and surgical staples, and the animals were allowed to recover before being returned to their home cage for 7 days before spinal *in vivo* photostimulation.

### In vivo photostimulation and behaviour

Animals were briefly anaesthetised (isoflurane, 5%) to attach a fiber optic patch cord (400 nm core, Thor Labs) to the implanted fiber optic probe before being placed in a small Perspex testing cylinder (10 cm diameter, 30 cm height) and allowed to habituate for 30 mins in the three days preceding photostimulation and for 20 minutes prior to photostimulation. The patch cord was attached to a high intensity LED light source (DC2100, 470 nm, Thor Labs) and photostimulation (10 mW, 10 ms pulses @ 10 Hz for 10 s) was delivered. Behavioural responses were recorded using a Panasonic video camera (Pansonic HC-V770M, Panasonic, Kadoma, Japan). In all experiments, animals were first introduced and acclimatised to the testing chamber for 3 days prior to testing, and video recordings captured 5 minutes of behaviour before and after photostimulation.

In some experiments the testing conditions were altered to address specific aspects of the photostimulation response. The relationship between stimulation intensity and behaviour was assessed in a subset of animals (n=6) received varying photostimulation intensities (0.5-20 mW, 10 ms pulses @ 10 Hz for 10 s), with a 30 min break between each stimulus. To assess the functional consequences of repeated spinal photostimulation, animals (n=9) received 2 photostimuli delivered 2 mins apart. To test the nociceptive nature of spinal photostimulation, animals (n=5) were administered morphine 30 minutes prior to photostimulation. Three morphine doses were assessed (3, 10 and 30mg/kg, s.c), as well as a saline vehicle control. Animals first underwent two photostimulation intensities (10 and 20 mW) with no morphine to determine baseline responses. Drug treatments were randomly assigned such that each animal received all concentrations and a 48 h interval between each drug administration allowed morphine washout. In experiments to assess the activation of higher order brain regions in response to photostimulation, animals (n=5 CR^cre^;Ai32, n=5 CReGFP) were placed in a testing chamber (30 cm length, 25 cm width and 40 cm height) with food and water available *ad libitum* for 6 hrs to eliminate any Fos activation caused by handling, the environment, or anaesthesia. Animals received photostimulation (10 mW, 10 ms pulses @ 10 Hz for 10 s) before being left for a further 2 h, to allow development of Fos expression, and then perfused transcardially with 4% PFA. Brains were dissected and post-fixed in 4% PFA overnight then stored in 30% sucrose. Serial sections were cut from the forebrain (40 µm) and brainstem (50 µm) using a freezing microtome (Leica Microsystems, SM2000R) and a 1 in 4 series were processed for Fos protein labelling (1:5000, rabbit polyclonal. Santa Cruz Biotechnology, CA, USA) as previously described ^50^. Fos positive cells were then manually counted from cingulate, insula, primary somatosensory cortex, parabrachial nucleus and periaqueductal grey. Counts were made on at least 4 sections ipsilateral to the stimulus side at 8.5X magnification.

In photo-inhibition experiments the fiber optic patch cord was attached to CR-NpHR animals (n = 8) which were then acclimatised to the testing tube as described previously. The patch cord was attached to the same laser described in *in vitro* methods. The simplified up down method (SUDO) (Bonin et al., 2014) of von Frey testing was used to establish mechanical withdrawal thresholds both with and without photo-inhibition. Animal were habituated in the testing chamber for 30 min for the 3 days prior to testing. Over four days of testing the von Frey threshold testing was assessed in each animal once with and once without photo-inhibition in alternating order. Withdrawal scores averaged across the four trial days and converted to withdrawal threshold in grams (Bonin et al., 2014).

All *in vivo* photostimulation-induced behaviour was analysed using JWatcher v1.0 event recorder (Blumstein and Daniel, 2007). Behavioural responses were encoded from the video recordings of photostimulation including 5 minutes pre- and post-photostimulation, played back at half-speed (30 fps). All behaviours targeted at the left or right hind limbs and the midline were coded. In a subset of videos coding was expanded to differentiate left/right paw and leg, as well as back and tail. The duration of all targeted behaviours was then binned (time epochs) and converted to a colour scale showing the proportion of epoch spent in specific behaviours for visualization.

### Conditioned place aversion testing

Conditioned place aversion (CPA) testing was used to assess the aversive nature of CR-ChR2 photostimulation. The CPA apparatus consisted of a two-chamber black perspex box (50 cm length, 25 cm width and 50 cm height) with a divider allowing free access to each chamber. To differentiate the two chambers one side contained cross-hatched markings using tape on the floor and crosses on the walls. On the first experimental day each animals baseline preference was determined. The optic patch cord was attached (as described above) and animal placed in the centre of the CPA apparatus. Animals were allowed to freely move between both chambers for 10 mins, prior to commencement of data collection. The chamber where an animal spent the most time was deemed the preferred side, and subsequently designated ‘photostimulation on’ while the non-preferred side was designated ‘photostimulation off’. Animals then underwent 4 × 20 min trials over 2 days (morning and afternoon), with the condition that entry into the ‘photostimulation on’ chamber triggered photostimulation (10 mW, 10 ms pulses @ 10 Hz for 10 s in every minute ie. 10 s on, 50 s off) until the animal returned to the ‘photostimulation off’ chamber. Following the CPA trials, the persistence of the CPA memory was assessed via short term (1 h post testing – STM) and long term tests (24 h post testing – LTM). In these tests animals were allowed to freely move around the CPA apparatus for 10 mins with no photostimulation. All trials and tests were captured from above the CPA apparatus (via the video camera), digitized, then analysed using semi-automated behavioural tracking procedures within Ethovision software (Noldus Information Technology, Wageningen, Netherlands).

### Statistical analysis

All data are presented as mean ± the standard error of the mean (SEM) unless otherwise stated. Shapiro-Wilk’s test determined if data were normally distributed. For normally distributed data one-way ANOVAs were performed with a student Newman-Keuls *post-hoc* test to compare oEPSC and oIPSC properties between neuron groups and for all behaviour analyses. Non-normally distributed data was compared using the Kruskal Wallis test with Wilcoxon–Mann–Whitney *post-hoc* testing. Paired t-tests compared sEPSC frequency before and after photostimulation in CR^+^ neuron and projection neuron populations.

## Acknowledgements

We thank Dr Philippe Ciofi for the guinea pig anti-somatostatin primary antibody, and both Christine Watt and Robert Kerr for expert technical assistance. This work was funded by the National Health and Medical Research Council (NHMRC) of Australia (grants 631000 and 1043933 to B.A.G, 1067146 to R.J.C. and 1125478 to C.V.D), the Biotechnology and Biological Sciences Research Council (BBSRC) of the United Kingdom (grant BB/J000620/1 and BB/P007996/1 to D.I.H.), and the Hunter Medical Research Institute (B.A.G. and R.J.C.).

## Author Contributions

K.M.S., R.J.C., P.J., C.V.D., D.I.H. and B.A.G. conceived and designed the research study; K.M.S., T.J.B, A.C. O.D., K.A.B., J.A.I., and S.A.D. conducted experiments and acquired data; M.W. kindly provided reagents; K.M.S., T.J.B, M.A.G, A.C., O.D., S.A.D., J.A.I., K.A.B., D.I.H., and B.A.G. analyzed data; K.M.S., C.V.D., D.I.H. and B.A.G. wrote the manuscript; all authors edited the final version of the manuscript.

## Competing financial interests

The authors do not have any conflict of interest.

---

**Supplementary Figure 1.**
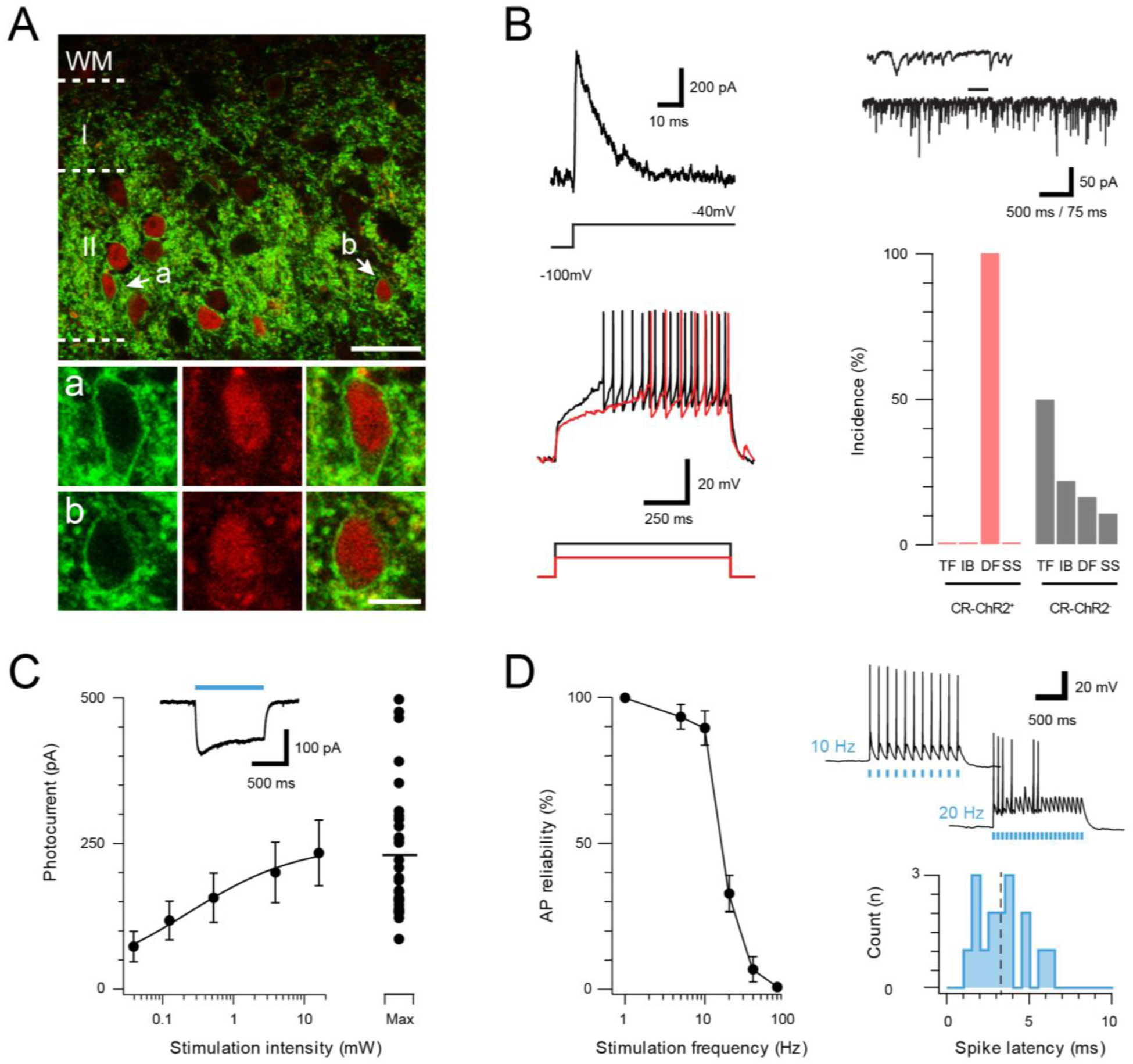
ChR2 expression in Excitatory CR^+^ neurons. (**A**), Upper panel compares ChR2YFP-IR (green) and CR-IR profiles (red). There is a high degree of colocalization in LII (71 ± 2% ChR2YFP-IR neurons express CR-IR, and 78 ± 4% of CR-IR neurons express ChR2YFP-IR). Lower panels show neurons denoted ‘a’ and ‘b’ from upper panel at high magnification; ChR2YFP-IR (left), CR-IR (right), merge (center), scale = 25μm (upper) and 5μm (lower). (**B**), Excitatory CR^+^ neurons exhibited several characteristic electrophysiological features including the voltage gated potassium current Ia (upper left, protocol below), high frequency spontaneous excitatory drive (upper right), and delayed firing (DF) discharge in response to depolarizing current injection (lower left, current step protocol below). Bar graph (lower, right) highlights the uniform incidence of DF-AP discharge in excitatory CR^+^ positive neurons (red) when compared to a random sample of CR negative neurons (grey) in the same region (TF = tonic firing, IB = Initial bursting, DF = Delayed firing, SS = Single spiking). (**C**), Plot shows relationship between photostimulation intensities (0.039-16 mW) and photocurrent amplitude, error bars = SEM. Note maximum photostimulation intensity (16 mW) shows photocurrent data for individual recordings. Inset, example photocurrent response with blue bar indicating photostimulus duration. (**D**), Plot (left) shows reliability of evoked AP discharge at various photostimulation frequencies. APs were reliably evoked by frequencies up to 10 Hz. Representative traces (upper, right) showing reliable responses at 10 Hz but not 20 Hz photostimulation. Histogram (lower, right) shows the distribution of recruitment latency (time between the onset of photostimulation and the AP response) for CR-ChR2 recordings (dashed line shows mean of 3.2 ms).

**Supplementary Figure 2.**
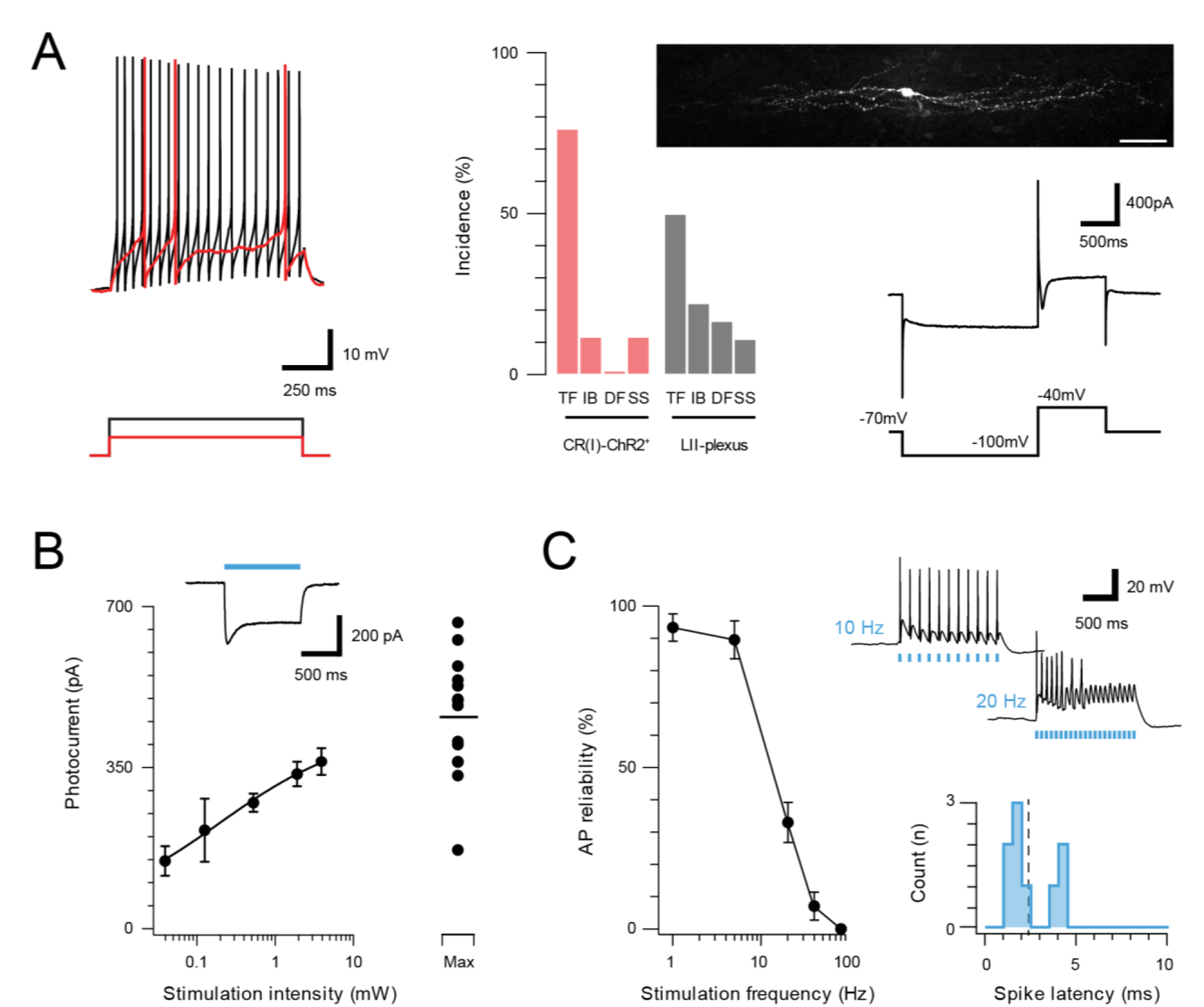
Inhibitory CR neurons express ChR2. A subset of inhibitory CR^+^ neurons, identified by extensive rostrocaudal processes, exhibited characteristic electrophysiological features described in previous work ^5^. (**A**) Most inhibitory CR^+^ cells responded with a tonic AP discharge (top left, protocol below). Bar graph (middle) shows elevated incidence of tonic discharge in inhibitory CR^+^ neurons when compared to a random sample of CR negative neurons (grey) from the same region (TF = tonic firing, IB = Initial bursting, DF = Delayed firing, SS = Single spiking). All neurobiotin-recovered inhibitory CR^+^ neurons exhibited islet like morphology (upper, right, scale = 20 μm) and expressed the I_h_ and I_Ca_ voltage activated currents (lower right, protocol below). (**B**), Group data for photocurrent amplitude at different light intensities (0.039-16 mW; error bars = SEM). Inset shows example photocurrent response for a 1s blue light stimulus, blue bar highlights photostimulus duration. Note, maximum photostimulation intensity (16 mW) shows photocurrent data for individual recordings. (**C**), Plot (left) shows reliability of evoked AP discharge at various photostimulation frequencies. APs were reliably evoked for stimulation frequencies of up to 10 Hz. Representative traces (upper, right) showing reliable responses at 10 Hz but not 20 Hz photostimulation. Histogram (lower, right) shows the distribution of recruitment latency (time between the onset of photostimulation and the AP response) for CR-ChR2^+^ recordings (dashed line = mean, 2.2 ms).

**Supplementary Figure 3.**
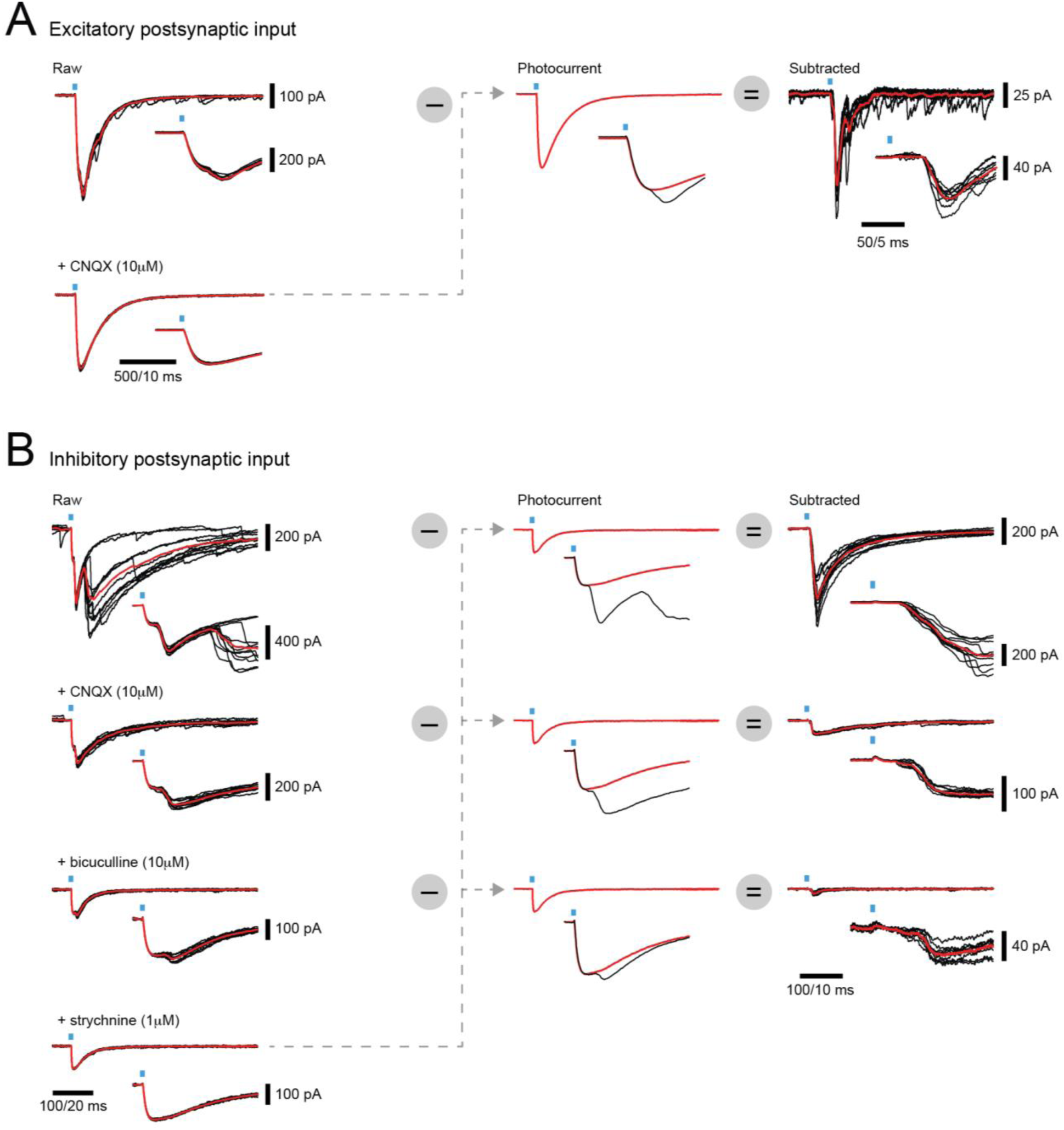
Isolation of synaptic responses in CR-ChR2^+^ neurons by photocurrent subtraction. (**A**), overlaid traces (left) show voltage clamp recordings of excitatory responses to 10 photostimulation sweeps (average in red) under baseline conditions (upper) and after bath addition of CNQX (10 μM). Insets show expanded response onset (blue bar highlights photostimulus). Note baseline response has two components at onset, the photocurrent and a synaptic current, but only the synaptic component is blocked by CNQX. The averaged photocurrent is isolated, rescaled to match the amplitude in individual baseline traces (middle), and then subtracted to yield isolated synaptic responses (right) to CR-ChR2 photostimulation. (**B**), overlayed traces arranged as in **A**, except voltage clamp recordings are inhibitory responses at baseline (upper), and after bath addition of CNQX (10 μM), bicuculline (10 μM), and strychnine (1 μM). Note multiple components at onset including both photocurrents and a synaptic current, but CNQX bicuculline, and strychnine are required to isolate the photocurrent. Photocurrent subtraction is then performed for each drug condition to yield the total inhibitory response with monosynaptic and polysynaptic components (upper), the monosynaptic inhibitory response (middle), and the glycinergic inhibitory response (lower) to CR-ChR2 photostimulation.

**Supplementary Figure 4.**
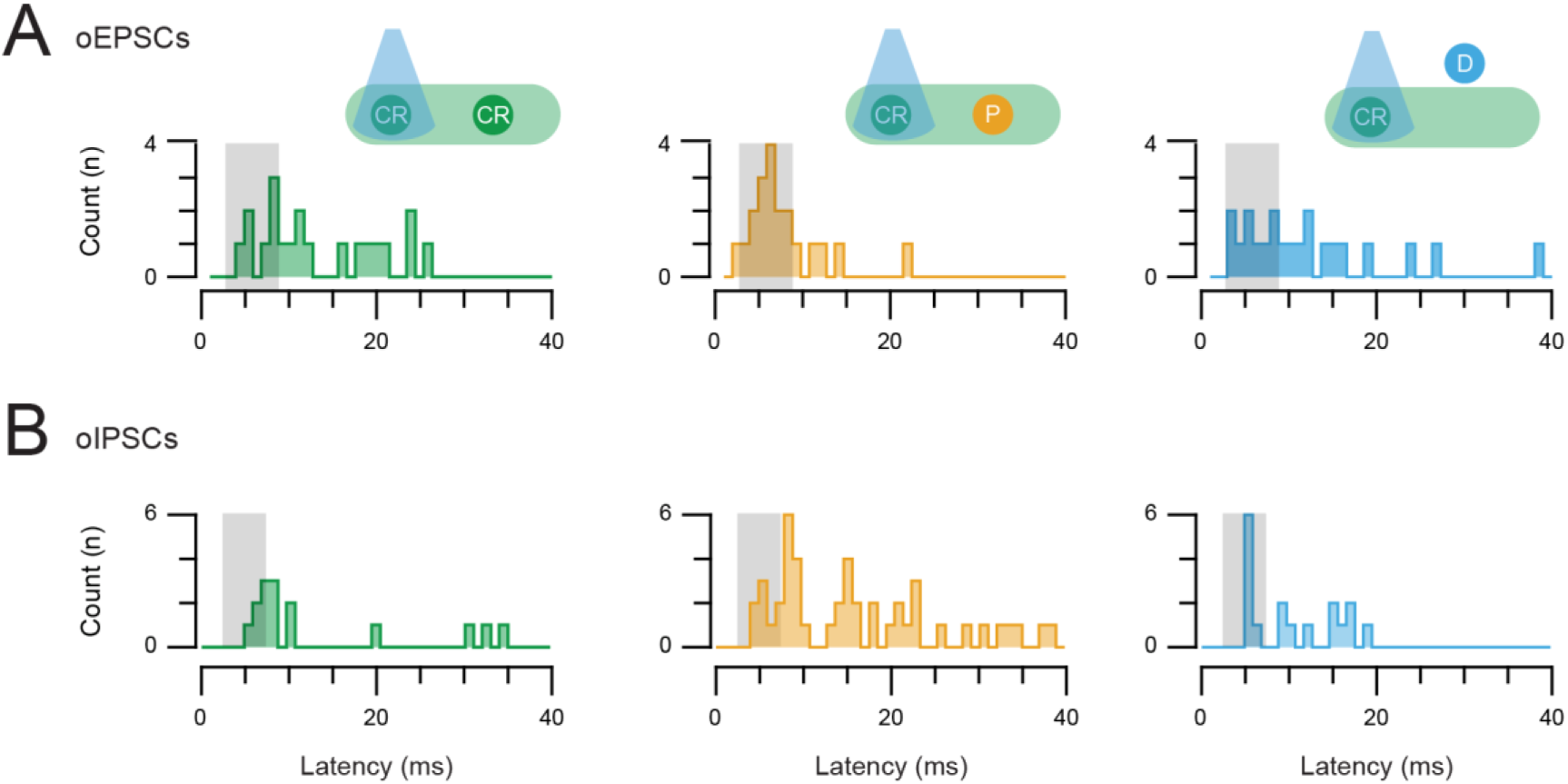
Photostimulation response latencies for oEPSC and oIPSC inputs. (**A-B**) The latency of all photostimulation response components (oEPSCs and oIPSCs, respectively) were measured and pooled for CR-ChR2 recordings, neurons within the CR^+^ plexus, and neurons dorsal to this region. Peristimulus histograms plot the onset latency of all synaptic responses components for each population (CR^+^, plexus, and dorsal). Grey box indicates the latency window for putative direct (monosynaptic) inputs. Note a proportion of responses in all populations exhibit latencies consistent with direct and indirect input.

**Supplementary Figure 5.**
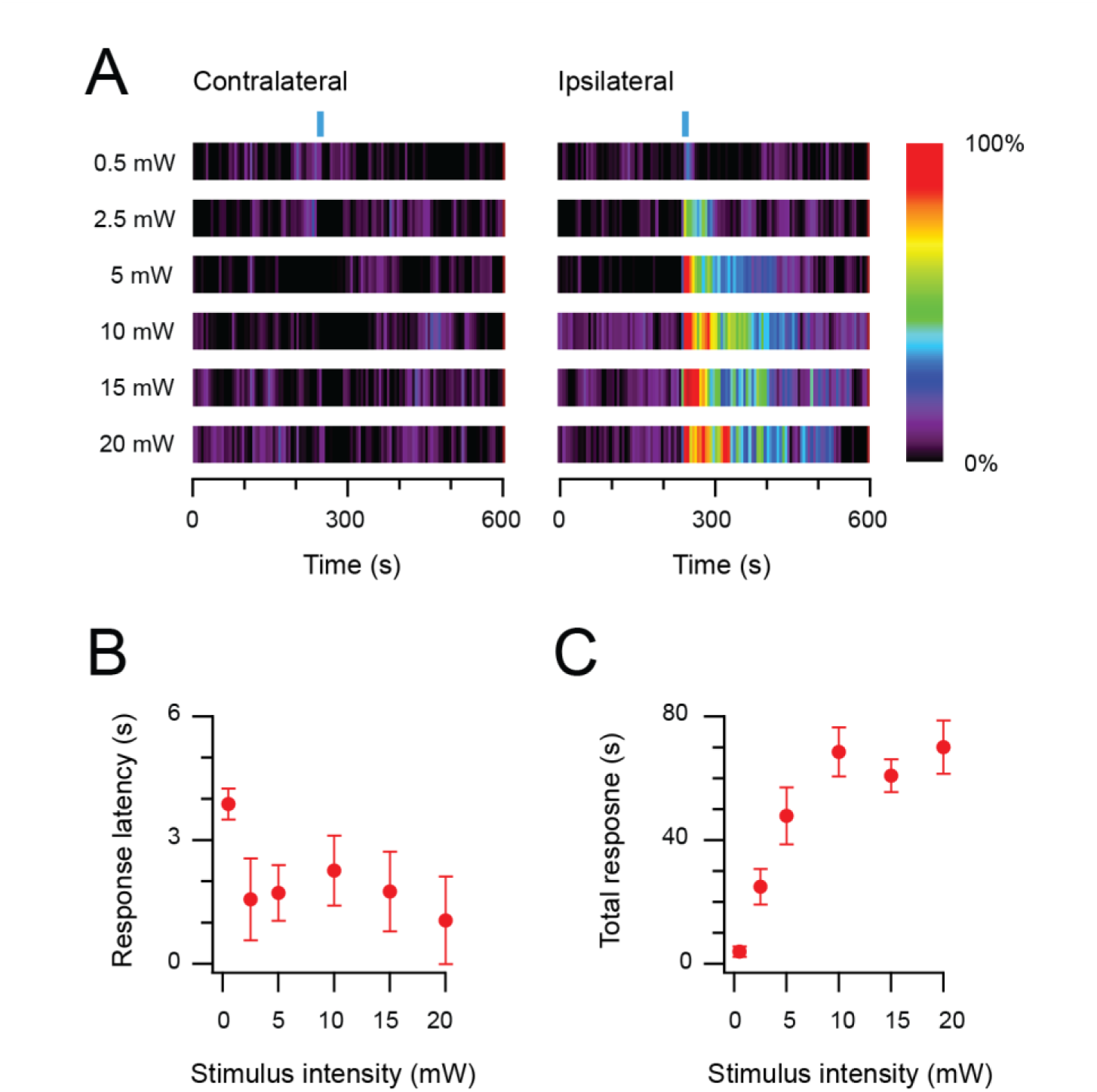
Photostimulation evokes an intensity dependent nociceptive behavioural response in CRcre;Ai32 mice. (**A**), Heat bars show averaged photostimulation (blue bar denotes photostimulation period) responses targeted to the contralateral hindlimb (left) and ipsilateral (right) hindlimb as photostimulus intensity is increased (0.5 – 20 mW, 10 Hz, 10 s). Responses are binned in 5 s epochs with bin color coding the percent time groomed (red = 100%, black = 0%). Reponses increased with photostimulation intensity and were focused to the ipsilateral hindlimb. (**B**), Plots compare response latency, and total response as stimulus intensity is increased. Note, at lower stimulus intensities some animals did not exhibit responses during the photostimulation period, and total response durations increased with photostimulation intensity before stabilizing between 10-15 mW.

**Supplementary Figure 6.**
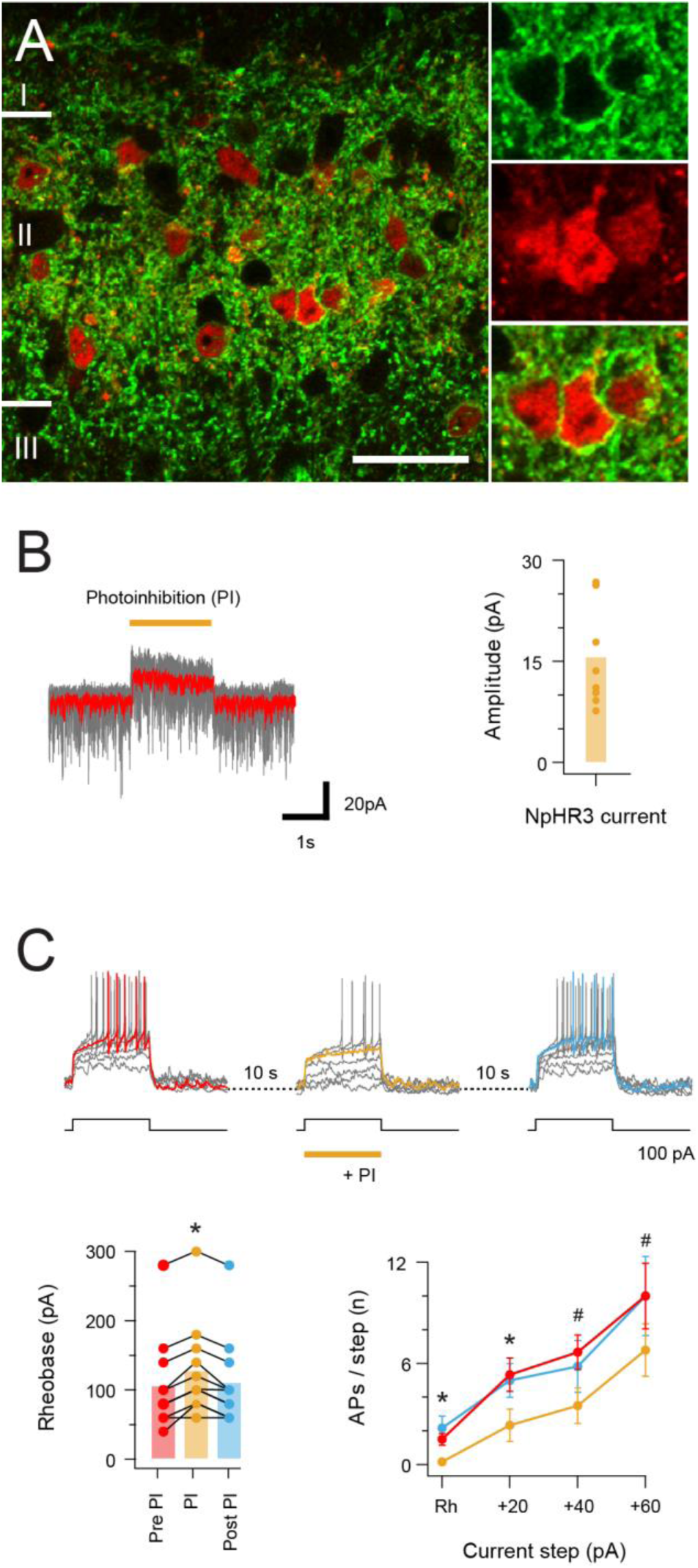
Halorhodopsin-mediated photoinhibition of CR^+^ neurons in CRcre;Ai32 mice. (**A**), Image panels (left) compare NpHR3YFP-IR (green) and CR-IR profiles (red). There is a high incidence of colocalisation of CR-IR in NpHR3YFP-IR neurons in laminae I and II (82.2%, ± 1.27), and of YFP expression in CR-IR neurons (94.1%, ± 4.27), Scale bar = 20 μm. (**B**), Traces (right) are overlaid recordings from a CR^+^ neuron during NpHR3-mediated photoinhibition (orange bar). The average response (red trace) highlight a stimulus-locked outward current resulting from NpHR3 activation. Plot (right) shows group data for NpHR3 current amplitude during photoinhibition. (**C**), Overlaid traces show CR^+^ neuron recording during 3 successive series of depolarizing current injection (20 pA increments, 800 ms duration) to activate AP discharge, separated by 10 s intervals. Sustained NpHR3-mediated photoinhibition was applied for the second current injection series. Photoinhibition suppressed current-evoked AP discharge, highlighted in group data plots comparing rheobase current (lower left) and current step versus AP number (lower right) during photostimulation with these properties pre- and post-photostimulation.

## References

Baba, H., Kohno, T., Moore, K. A. & Woolf, C. J. 2001. Direct activation of rat spinal dorsal horn neurons by prostaglandin E2. J Neurosci, 21, 1750–6.

Basbaum, A. I., Bautista, D. M., Scherrer, G. & Julius, D. 2009. Cellular and molecular mechanisms of pain. Cell, 139, 267–84.

Blumstein, D. T. & Daniel, J. C. 2007. Quantifying behaviour the JWatcher way, Sinauer Associates Inc.

Bonin, R. P., Bories, C. & De Koninck, Y. 2014. A simplified up-down method (SUDO) for measuring mechanical nociception in rodents using von Frey filaments. Mol Pain, 10, 26.

Bonin, R. P., Wang, F., Desrochers-Couture, M., Ga Secka, A., Boulanger, M. E., Cote, D. C. & De Koninck, Y. 2016. Epidural optogenetics for controlled analgesia. Mol Pain, 12.

Braz, J. M., Juarez-Salinas, D., Ross, S. E. & Basbaum, A. I. 2014. Transplant restoration of spinal cord inhibitory controls ameliorates neuropathic itch. J Clin Invest, 124, 3612–6.

Caputi, A., Rozov, A., Blatow, M. & Monyer, H. 2009. Two calretinin-positive GABAergic cell types in layer 2/3 of the mouse neocortex provide different forms of inhibition. Cereb Cortex, 19, 1345–59.

Christensen, A. J., Iyer, S. M., Francois, A., Vyas, S., Ramakrishnan, C., Vesuna, S., Deisseroth, K., Scherrer, G. & Delp, S. L. 2016. In Vivo Interrogation of Spinal Mechanosensory Circuits. Cell Rep, 17, 1699–1710.

Ciofi, P., Leroy, D. & Tramu, G. 2006. Sexual dimorphism in the organization of the rat hypothalamic infundibular area. Neuroscience, 141, 1731–45.

Coull, J. A., Boudreau, D., Bachand, K., Prescott, S. A., Nault, F., Sik, A., De Koninck, P. & De Koninck, Y. 2003. Trans-synaptic shift in anion gradient in spinal lamina I neurons as a mechanism of neuropathic pain. Nature, 424, 938–42.

Cui, L., Miao, X., Liang, L., Abdus-Saboor, I., Olson, W., Fleming, Michael S., Ma, M., Tao, Y.-X. & Luo, W. 2016. Identification of Early RET+ Deep Dorsal Spinal Cord Interneurons in Gating Pain. Neuron, 91, 1137–1153.

Duan, B., Cheng, L., Bourane, S., Britz, O., Padilla, C., Garcia-Campmany, L., Krashes, M., Knowlton, W., Velasquez, T., Ren, X., Ross, S., Lowell, B. B., Wang, Y., Goulding, M. & Ma, Q. 2014. Identification of spinal circuits transmitting and gating mechanical pain. Cell, 159, 1417–1432.

Foster, E., Wildner, H., Tudeau, L., Haueter, S., Ralvenius, William T., Jegen, M., Johannssen, H., Hösli, L., Haenraets, K., Ghanem, A., Conzelmann, K.-K., Bösl, M. & Zeilhofer, Hanns U. 2015. Targeted Ablation, Silencing, and Activation Establish Glycinergic Dorsal Horn Neurons as Key Components of a Spinal Gate for Pain and Itch. Neuron, 85, 1289–1304.

Francois, A., Low, S. A., Sypek, E. I., Christensen, A. J., Sotoudeh, C., Beier, K. T., Ramakrishnan, C., Ritola, K. D., Sharif-Naeini, R., Deisseroth, K., Delp, S. L., Malenka, R. C., Luo, L., Hantman, A. W. & Scherrer, G. 2017. A Brainstem-Spinal Cord Inhibitory Circuit for Mechanical Pain Modulation by GABA and Enkephalins. Neuron, 93, 822–839.e6.

Ganley, R. P., Iwagaki, N., Del Rio, P., Baseer, N. & Dickie, A. C. 2015. Inhibitory Interneurons That Express GFP in the PrP-GFP Mouse Spinal Cord Are Morphologically Heterogeneous, Innervated by Several Classes of Primary Afferent and Include Lamina I Projection Neurons among Their Postsynaptic Targets. 35, 7626–42.

Graham, B. A., Brichta, A. M. & Callister, R. J. 2004. In vivo responses of mouse superficial dorsal horn neurones to both current injection and peripheral cutaneous stimulation. J Physiol, 561, 749–63.

Graham, B. A., Brichta, A. M., Schofield, P. R. & Callister, R. J. 2007. Altered potassium channel function in the superficial dorsal horn of the spastic mouse. J Physiol, 584, 121–36.

Graham, B. A., Schofield, P. R., Sah, P. & Callister, R. J. 2003. Altered inhibitory synaptic transmission in superficial dorsal horn neurones in spastic and oscillator mice. J Physiol, 551, 905–16.

Graham, B. A., Tadros, M. A., Schofield, P. R. & Callister, R. J. 2011. Probing glycine receptor stoichiometry in superficial dorsal horn neurones using the spasmodic mouse. J Physiol, 589, 2459–74.

Gutierrez-Mecinas, M., Bell, A. M., Marin, A., Taylor, R., Boyle, K. A., Furuta, T., Watanabe, M., Polgar, E. & Todd, A. J. 2017. Preprotachykinin A is expressed by a distinct population of excitatory neurons in the mouse superficial spinal dorsal horn including cells that respond to noxious and pruritic stimuli. Pain, 158, 440–456.

Gutierrez-Mecinas, M., Furuta, T., Watanabe, M. & Todd, A. J. 2016. A quantitative study of neurochemically defined excitatory interneuron populations in laminae I-III of the mouse spinal cord. Mol Pain, 12.

Hachisuka, J., Omori, Y., Chiang, M. C., Gold, M. S., Koerber, H. R. & Ross, S. E. 2018. Wind-up in lamina I spinoparabrachial neurons: a role for reverberatory circuits. Pain, 159, 1484–1493.

Hrabovszky, E., Csapo, A. K., Kallo, I., Wilheim, T., Turi, G. F. & Liposits, Z. 2006. Localization and osmotic regulation of vesicular glutamate transporter-2 in magnocellular neurons of the rat hypothalamus. Neurochem Int, 48, 753–61.

Hughes, D. I., Boyle, K. A., Kinnon, C. M., Bilsland, C., Quayle, J. A., Callister, R. J. & Graham, B. A. 2013. HCN4 subunit expression in fast-spiking interneurons of the rat spinal cord and hippocampus. Neuroscience, 237, 7–18.

Kato, G., Kosugi, M., Mizuno, M. & Strassman, A. M. 2011. Separate inhibitory and excitatory components underlying receptive field organization in superficial medullary dorsal horn neurons. J Neurosci, 31, 17300–5.

Lu, Y., Dong, H., Gao, Y., Gong, Y., Ren, Y., Gu, N., Zhou, S., Xia, N., Sun, Y.-Y., Ji, R.-R. & Xiong, L. 2013. A feed-forward spinal cord glycinergic neural circuit gates mechanical allodynia. The Journal of Clinical Investigation, 123, 4050–4062.

Lu, Y. & Perl, E. R. 2003. A specific inhibitory pathway between substantia gelatinosa neurons receiving direct C-fiber input. J Neurosci, 23, 8752–8.

Lu, Y. & Perl, E. R. 2005. Modular organization of excitatory circuits between neurons of the spinal superficial dorsal horn (laminae I and II). J Neurosci, 25, 3900–7.

Lui, F. & Ng, K. F. 2011. Adjuvant analgesics in acute pain. Expert Opin Pharmacother, 12, 363–85.

Luo, C., Kuner, T. & Kuner, R. 2014. Synaptic plasticity in pathological pain. Trends Neurosci, 37, 343–55.

Meyer, A. H., Katona, I., Blatow, M., Rozov, A. & Monyer, H. 2002. In vivo labeling of parvalbumin-positive interneurons and analysis of electrical coupling in identified neurons. J Neurosci, 22, 7055–64.

Miraucourt, L. S., Dallel, R. & Voisin, D. L. 2007. Glycine inhibitory dysfunction turns touch into pain through PKCgamma interneurons. PLoS One, 2, e1116.

Moore, K. A., Kohno, T., Karchewski, L. A., Scholz, J., Baba, H. & Woolf, C. J. 2002. Partial peripheral nerve injury promotes a selective loss of GABAergic inhibition in the superficial dorsal horn of the spinal cord. J Neurosci, 22, 6724–31.

Nakamura, M., Sato, K., Fukaya, M., Araishi, K., Aiba, A., Kano, M. & Watanabe, M. 2004. Signaling complex formation of phospholipase Cbeta4 with metabotropic glutamate receptor type 1alpha and 1,4,5-trisphosphate receptor at the perisynapse and endoplasmic reticulum in the mouse brain. Eur J Neurosci, 20, 2929–44.

Neumann, S., Braz, J. M., Skinner, K., Llewellyn-Smith, I. J. & Basbaum, A. I. 2008. Innocuous, not noxious, input activates PKCgamma interneurons of the spinal dorsal horn via myelinated afferent fibers. J Neurosci, 28, 7936–44.

Paxinos, G. & Franklin, K. 2001. The mouse brain in stereotaxic coordinates (2nd edition), Academic Press.

Peirs, C. & Seal, R. P. 2016. Neural circuits for pain: Recent advances and current views. Science, 354, 578–584.

Peirs, C., Williams, S.-P. G., Zhao, X., Walsh, C. E., Gedeon, J. Y., Cagle, N. E., Goldring, A. C., Hioki, H., Liu, Z., Marell, P. S. & Seal, R. P. 2015. Dorsal Horn Circuits for Persistent Mechanical Pain. Neuron, 87, 797–812.

Petitjean, H., Pawlowski, Sophie A., Fraine, Steven L., Sharif, B., Hamad, D., Fatima, T., Berg, J., Brown, Claire M., Jan, L.-Y., Ribeiro-Da-Silva, A., Braz, Joao M., Basbaum, Allan I. & Sharif-Naeini, R. 2015. Dorsal Horn Parvalbumin Neurons Are Gate-Keepers of Touch-Evoked Pain after Nerve Injury. Cell Reports, 13, 1246–1257.

Polgar, E., Durrieux, C., Hughes, D. I. & Todd, A. J. 2013. A quantitative study of inhibitory interneurons in laminae I-III of the mouse spinal dorsal horn. PLoS One, 8, e78309.

Price, T. J. & Prescott, S. A. 2015. Inhibitory regulation of the pain gate and how its failure causes pathological pain. Pain, 156, 789–92.

Proudlock, F., Spike, R. C. & Todd, A. J. 1993. Immunocytochemical study of somatostatin, neurotensin, Gaba, and glycine in rat spinal dorsal horn. J Comp Neurol, 327, 289–97.

Ptak, K., Burnet, H., Blanchi, B., Sieweke, M., De Felipe, C., Hunt, S. P., Monteau, R. & Hilaire, G. 2002. The murine neurokinin NK1 receptor gene contributes to the adult hypoxic facilitation of ventilation. Eur J Neurosci, 16, 2245–52.

Punnakkal, P., Von Schoultz, C., Haenraets, K., Wildner, H. & Zeilhofer, H. U. 2014. Morphological, biophysical and synaptic properties of glutamatergic neurons of the mouse spinal dorsal horn. J Physiol, 592, 759–76.

Ross, S. E., Mardinly, A. R., Mccord, A. E., Zurawski, J., Cohen, S., Jung, C., Hu, L., Mok, S. I., Shah, A., Savner, E., Tolias, C., Corfas, R., Chen, S., Inquimbert, P., Xu, Y., Mcinnes, R. R., Rice, F. L., Corfas, G., Ma, Q., Woolf, C. J. & Greenberg, M. E. 2010. Loss of inhibitory interneurons in the dorsal spinal cord and elevated itch in Bhlhb5 mutant mice. Neuron, 65, 886–898.

Santos, S. F., Rebelo, S., Derkach, V. A. & Safronov, B. V. 2007. Excitatory interneurons dominate sensory processing in the spinal substantia gelatinosa of rat. J Physiol, 581, 241–54.

Schiffmann, S. N., Cheron, G., Lohof, A., D’Alcantara, P., Meyer, M., Parmentier, M. & Schurmans, S. 1999. Impaired motor coordination and Purkinje cell excitability in mice lacking calretinin. Proc Natl Acad Sci U S A, 96, 5257–62.

Schwarz, L. A., Miyamichi, K., Gao, X. J., Beier, K. T., Weissbourd, B., Deloach, K. E., Ren, J., Ibanes, S., Malenka, R. C., Kremer, E. J. & Luo, L. 2015. Viral-genetic tracing of the input-output organization of a central noradrenaline circuit. Nature, 524, 88–92.

Smith, K. M., Boyle, K. A., Madden, J. F., Dickinson, S. A., Jobling, P., Callister, R. J., Hughes, D. I. & Graham, B. A. 2015. Functional heterogeneity of calretinin-expressing neurons in the mouse superficial dorsal horn: implications for spinal pain processing. J Physiol, 593, 4319–39.

Smith, K. M., Boyle, K. A., Mustapa, M., Jobling, P., Callister, R. J., Hughes, D. I. & Graham, B. A. 2016. Distinct forms of synaptic inhibition and neuromodulation regulate calretinin-positive neuron excitability in the spinal cord dorsal horn. Neuroscience, 326, 10–21.

Takazawa, T. & Macdermott, A. B. 2010. Synaptic pathways and inhibitory gates in the spinal cord dorsal horn. Ann N Y Acad Sci, 1198, 153–8.

Tamas, G., Buhl, E. H., Lorincz, A. & Somogyi, P. 2000. Proximally targeted GABAergic synapses and gap junctions synchronize cortical interneurons. Nat Neurosci, 3, 366–71.

Todd, A. J. 2010. Neuronal circuitry for pain processing in the dorsal horn. Nat Rev Neurosci, 11, 823–36.

Torsney, C. & Macdermott, A. B. 2006. Disinhibition opens the gate to pathological pain signaling in superficial neurokinin 1 receptor-expressing neurons in rat spinal cord. J Neurosci, 26, 1833–43.

Woodruff, A. R. & Sah, P. 2007. Inhibition and synchronization of basal amygdala principal neuron spiking by parvalbumin-positive interneurons. J Neurophysiol, 98, 2956–61.

Woolf, C. J. & Fitzgerald, M. 1983. The properties of neurones recorded in the superficial dorsal horn of the rat spinal cord. J Comp Neurol, 221, 313–28.

Yu, F., Zhao, Z.-Y., He, T., Yu, Y.-Q., Li, Z. & Chen, J. 2017. Temporal and spatial dynamics of peripheral afferent-evoked activity in the dorsal horn recorded in rat spinal cord slices. Brain Research Bulletin, 131, 183–191.

Zeilhofer, H. U. 2005. The glycinergic control of spinal pain processing. Cell Mol Life Sci, 62, 2027–35.

Zeilhofer, H. U., Wildner, H. & Yevenes, G. E. 2012. Fast synaptic inhibition in spinal sensory processing and pain control. Physiol Rev, 92, 193–235.

